# Quantitative description of the phase separation behavior of the multivalent SLP65-CIN85 complex

**DOI:** 10.1101/2023.07.31.551239

**Authors:** Joachim Maier, Daniel Sieme, Leo E. Wong, Furqan Dar, Jürgen Wienands, Stefan Becker, Christian Griesinger

## Abstract

Biomolecular condensates play a major role in cell compartmentalization, besides membrane-enclosed organelles. The multivalent SLP65 and CIN85 proteins are downstream B cell receptor (BCR)-signaling effectors, required for a proper immune response. Both proteins phase separate together with vesicles to form pre-signaling clusters. Within this tripartite system, six PRMs of SLP65 interact promiscuously with three SH3 domains of the CIN85 monomer, establishing 18 individual SH3-PRM interactions whose individual dissociation constants we determined. Based on these 18 dissociation constants, we measured the phase separation properties of the natural SLP65/CIN85 system as well as designer constructs that emphasize the strongest SH3/PRM interactions. By modelling these various SLP65/CIN85 constructs with the program LASSI (LAttice simulation engine for Sticker and Spacer Interactions) we reproduced the observed phase separation properties. In addition, LASSI revealed a deviation in the experimental measurement, which was independently identified as a previously unknown intramolecular interaction. Thus, thermodynamic properties of the individual PRM/SH3 interactions allow to model the phase separation behavior of the SLP65/CIN85 system faithfully.

## Introduction

Liquid-liquid phase separation of biomolecules has emerged as a general principle for cellular organization [1, 2]. A diverse set of biomolecules can participate in phase separation, e.g. intrinsically disordered proteins [3-7], proteins with low-complexity segments [8], RNA [9] or RNA-binding proteins [10], multivalent proteins [11] as well as synaptic vesicles [12]. Phase separation has been found in cellular signaling pathways such as BCR and T cell receptor signaling [13-16]. The Src homology 2 domain-containing leukocyte protein of 65 kDa (SLP65) [17], also referred to as B-cell linker protein (BLNK) [18], is a signaling protein in the BCR signaling pathway. SLP65 and its cellular binding partner, Cbl-interacting protein of 85 kDa (CIN85) [19] are both multivalent proteins. They have an essential immunological function and deficiency of CIN85 or SLP65 leads to a compromised IgG response during bacterial infections [20-22]. In resting B cells, CIN85 and vesicle-associated SLP65 condense into pre-signaling clusters, which prime B cell responsiveness [14]. The assembly of these phase-separated pre-signaling clusters is driven by the binding of SLP65’s N-terminus to cytosolic vesicles [19], as well as the promiscuous interaction of SLP65’s PRMs with the three SH3 domains of CIN85 [14] and the trimerization of CIN85 by its CC domain [23] (Fig. 1*A*).

**Fig. 1:**
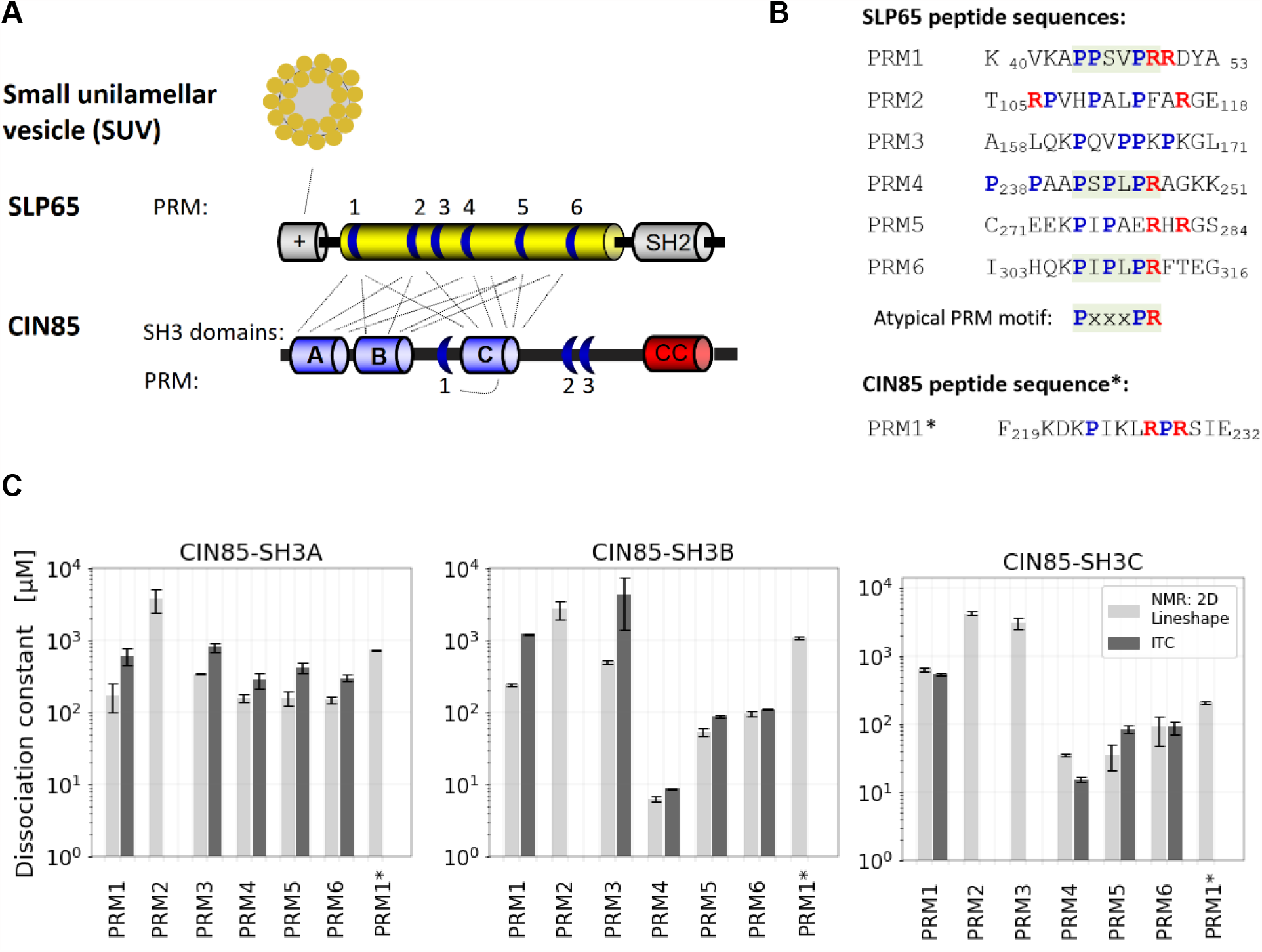
A) Schematic illustration of the interactions mediating tripartite phase-separation of SLP65, CIN85 and SUVs. SLP65’s positively charged N-terminus interacts with small unilamellar vesicles [14], while the C-terminal SH2 domain has a role in signal transduction (2002 Kabak). SLP65’s intrinsically disordered region (yellow) harbors six PRMs (blue regions), which promiscuously interact with the SH3A, -B and -C domains of CIN85 [14]. CIN85’s coiled-coil domain (CC) forms a trimer (not shown). Additionally, CIN85’s PRM1 interacts with SH3C (D. Sieme, BioRxiv). B) Overview of the sequences of the synthesized 14 amino acid long peptides derived from PRM regions. The atypical motif PxxxPR is highlighted in grey; Arg and Pro are colored red and blue, respectively. PRM1*, PRM1 of CIN85. C) Dissociation constants of single PRM-representing peptides to CIN85’s SH3A, SH3B and SH3C domains. K_D_ values for PRM1* were provided by D. Sieme (Sieme et al., BioRxiv). The 21 dissociation constants (K_D_s) of the SH3-PRM interactions range from 6 μM to _∼_4 mM. K_D_s were obtained by NMR (TITAN v1.5-3-g3566 [37], grey bars) and ITC titrations (black bars). Both methods show that PRM4 has the largest and second largest K_D_ in the interaction with SH3B and SH3C, respectively. No heat release was measured for the weak interactions of PRM2 with any SH3 domain and PRM3 with SH3C, thus no K_D_ is shown here.

The PRM_m_-SH3_n_ system is a prominent example for phase separation of multivalent proteins. Pioneering biophysical and computational studies worked with identical PRM and SH3 domains [24], yet, in SLP65 and CIN85 the 6 PRMs and the 3 SH3 domains differ in sequence (Fig. 1B). Therefore, the individual dissociation constants are expected to vary. Our approach is to relate the phase separation properties to individual modules of SLP65 and CIN85. As a first step, the dissociation constants were measured. In the next step, the phase separation was modelled based on the ranking of the underlying SH3-PRM interactions by using the open source program LASSI [25]. The program is geared towards multivalent proteins by implementing a lattice based, coarse-grained Monte Carlo simulation. Previous phase separation models of LASSI correlated phase separation properties with the interaction strengths of individual amino acids with each other implemented as an energy matrix [26]. In the herein applied LASSI model, we interpreted each SH3 domain and each PRM as a single sticker and calibrated the linker lengths with the diameter of the SH3 domains. Isothermal phase diagrams were constructed by running a set of simulations of the binary mixtures, setting up different combinations of SLP65 and CIN85 constructs and their mutants. The constructs were also investigated experimentally, revealing a good correlation of the experimentally measured and simulated critical concentrations. We arrive at a descriptive LASSI model for the CIN85-SLP65 system, reproducing the experimental observations quantitatively.

## Results and Discussion

### Disentanglement of the promiscuous SH3-PRM interactions reveals the particular strong interaction of SLP65-PRM4 with CIN85’s SH3B domain

In order to elucidate the thermodynamics of phase separation upon SLP65 and CIN85 pre-signaling cluster formation, the dissociation constants (K_D_s) of the underlying SH3-PRM interactions were measured. The 18 known interactions stemming from the three SH3 domains of CIN85 SH3A, SH3B, SH3C and the six PRMs of SLP65 [Fig. S8 in Wong *et al*. [14]] were investigated by titrations. The K_D_s of the binary interactions were obtained from concentration dependent chemical shift changes by NMR spectroscopy (Fig. 1*C*; Fig. S1 and S2). Among the 18 titration experiments, the NMR resonances appeared in fast, slow and intermediate exchange regimes (Fig. S3). For illustration, the NMR spectra of the titration of SLP65-PRM6 to the ^15^N-SH3A shows most peaks in fast exchange, except of D16 and N51, which exhibit intermediate exchange (Fig. S3*A*, marked by arrows). An illustrative example for intermediate exchange is the titration of SLP65-PRM4 to ^15^N-SH3C (Fig. S3*B*). Slow exchange was observed in the titration of SLP65-PRM4 to ^15^N-SH3B (Fig. S3*C*). Moreover, the K_D_s were determined by isothermal titration calorimetry (ITC) (Fig. S4, upper panel). The measurements by ITC, as an independent technique, corroborated the K_D_s measured by NMR (Fig. 1*C*, Fig. S5). Both methods, ITC and NMR, demonstrate that PRM4 is the strongest binding motif, followed in rank by PRM5 and PRM6, while PRM1 and PRM3 interact weaker. PRM2 was found to be the weakest binding motif, for which no heat release could be measured by ITC (Fig. S4, lower panel).

### The strong-binding CIN85-BBB, but not the SLP65-3xPRM4 construct reduces the critical concentration for tripartite phase separation

Pre-signaling clusters composed of vesicle-associated SLP65 and CIN85 can be reconstituted *in vitro* by mixing recombinant SLP65, CIN85 and small unilamellar vesicles (SUVs) [14]. In order to investigate how the dissociation constant influences phase separation, weak- and strong-binding SLP65 and CIN85 constructs were designed (Fig. 2A) taking advantage of the previously determined dissociation constants (Fig. 1*C*). Specifically, we studied the effect of enhancing the interaction between SPL65 and CIN85 by introducing more or less of the strong PRM4/SH3B interactions. In one implementation with enhanced PRM4/SH3B interactions, the low-affine PRM5 and PRM6 were replaced by PRM4 leading to the SLP65-3xPRM4 construct. The second implementation to enhance the PRM4/SH3B interactions was the CIN85-3xSH3B construct, herein referred to as CIN85-BBB, in which each of the weaker binding SH3A and SH3C domains were replaced by the SH3B domain. On the other hand, to weaken the influence of the PRM4/SH3B interaction we used the SLP65-R247A construct that has a SH3 domain binding incompetent PRM4 due to the R247A mutation [27]. Due to higher yield in recombinant protein production, the C-terminally shortened versions SLP65_1-330_ and CIN85_1-333_ were produced and used for measurements. CIN85_1-333_ contains the SH3A, -B and -C domains and is herein referred to as CIN85-ABC construct. We note that the CIN85-ABC construct does not trimerize, in contrast to full length CIN85, which contains the C-terminal coiled-coil domain.

**Fig. 2:**
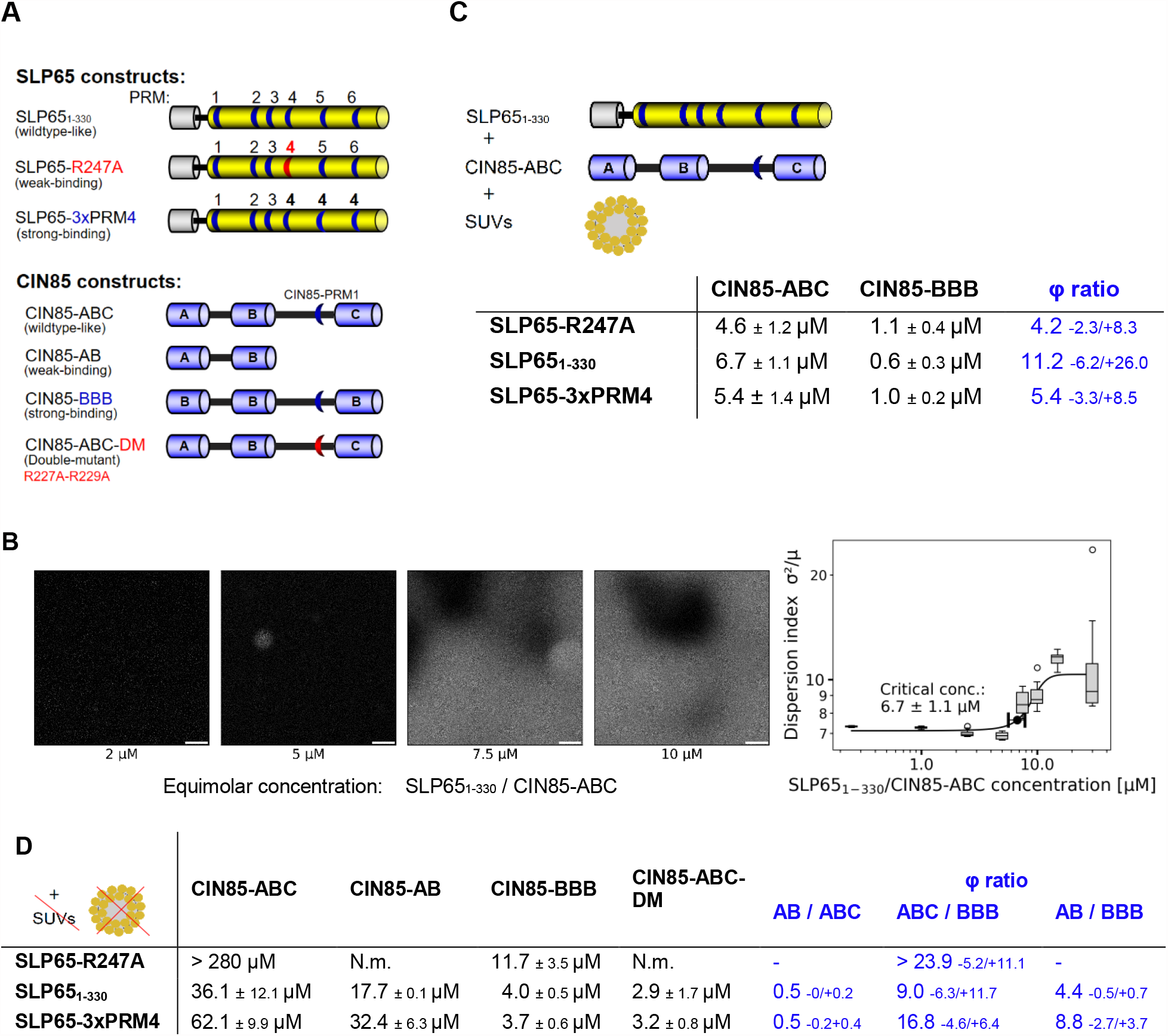
Exploring the phase separation behavior of the SH3-PRM system of CIN85 and SLP65. A) Construct design for weak- and strong-binding versions of SLP65 and CIN85. The SLP65_1-330_ construct, shortened at the C-terminus comprising residues 1-330, lacks the C-terminal SH2 domain, which is irrelevant for the CIN85 interaction. The weak-binding SLP65-R247A construct has a reduced affinity to CIN85 due to the R247A mutation, which deactivates the binding of PRM4. The strong-binding SLP65-3xPRM4 construct is designed by replacing both PRM5 and PRM6 by PRM4. The CIN85-ABC comprises residues 1-333. The weak-binding CIN85-AB construct contains only SH3A and SH3B. In the strong-binding CIN85-BBB construct, the SH3A and SH3C domain are each replaced by the strong binding SH3B domain. In the CIN85-ABC-double-mutant construct (DM) the CIN85-PRM1 is deactivated by the R227A and R229A mutations. B) Experimental determination of the critical concentration of phase separation: atto430LS-SLP65, CIN85 and SUV were mixed in a concentration series and imaged by confocal fluorescence microscopy. The critical concentration is obtained by a sigmoidal fit of the dispersion index (ratio of the variance σ to the mean μ). C) Critical concentrations of phase separation (φ_exp_) of mixture compositions of SUV + CIN85 + SLP65 mixed at equimolar protein concentrations. The strong binding version CIN85-BBB reduces φ_exp_ compared to the *wildtype*-like CIN85-ABC construct by a factor of 4 to 11, referred here as φ-ratio, whereas the weak- and strong-binding SLP65 versions, SLP65-R247A and SLP65-3xPRM4, respectively, have no effect on the critical concentration for phase separation. The φ-ratio is given as the number and the confidence interval is given towards smaller and larger numbers. For example: 4.2 – 2.3/+8.3 means that the confidence interval is from 4.2-2.3 = 1.9 to 4.2 + 8.3 = 12.5.D) Critical concentrations of phase separation φ_exp_ of SLP65/CIN85 mixtures without SUVs. φ ratios are calculated as the ratio of the critical concentrations *(continued legend Fig. 2)* of two different mixtures. φ_exp_ of the SLP65-R247A & CIN85-ABC mixture could not be fitted in our measurements with an upper end of the concentration series of 320 μM (Fig. S9), N.m. (not measured). The error for φ ratios is given as confidence interval as in Fig. 1C.

To elucidate the strength of binding of the described constructs, we measured their dissociation constants by ITC (Table S1). *Wildtype* SLP65_1-330_ and CIN85-ABC had a K_D_ of 1.36 ± 0.17 μM (Fig. S6*A*) which dropped by a factor of almost 6 (K_D_ = 0.24 ± 0.01 μM) when SLP65_1-330_ was replaced by SLP65-3xPRM4 (Fig. S6*B*). As expected, a larger K_D_ of 7.04 ± 1.12 μM was measured when SLP65_1-330_ was replaced by SLP65-R247A (Fig. S6*C*). The K_D_s of the interactions of the CIN85-BBB construct with either the SLP65_1-330_ or the SLP65-3xPRM4 construct were 0.7 ± 0.25 and 0.14 ± 0.06 μM, respectively (Fig. S6*D* and *E*) amounting to a change in K_D_s of around 2 when CIN85-ABC was replaced by CIN85-BBB (Table S1). The interaction of CIN85-BBB with SLP65-R247A was found similar in strength as CIN85-ABC with SLP65-R247A (Fig. S6*C* and *F*).

Subsequently, we explored the affinity-dependent phase separation behavior of the tripartite system by confocal fluorescence microscopy, imaging mixtures of the described constructs of SLP65 and CIN85 at equimolar concentration with vesicles (Fig. 2*B*). Since in solution, SLP65/CIN85/vesicle condensates sink down just after mixing, the visualization of condensates on the bottom of the microscopy slide was preferred over turbidity measurements in a vial [28]. The critical concentration of phase separation (φ_exp_) was determined by fitting the dispersion index (Fig. S7 and S8), since it is a simpler method than dual-color fluorescence cross-correlation spectroscopy [29] and since it is a more objective measure than visual evaluation [30]. For the measurements, SLP65 partially labeled with fluorescein (up to 10 μM) was used. The mixture of SLP65_1-330_, CIN85-ABC and SUVs resulted in φ_exp_ of 6.7 ± 1.1 μM (Fig. 2*C*; Fig. S7A), while the weak-binding version SLP65-R247A mixed with CIN85-ABC and SUVs resulted in a lower φ_exp_ (4.5 ± 1.2 μM, Fig. S7*B*). The strong-binding version SLP65-3xPRM4 with CIN85-ABC and SUVs delivered a φ_exp_ of 5.5 ± 1.4 μM (Fig. S7*C*). Thus, inactivating PRM4 (SLP65-R247A) or increasing its number (SLP65-3xPRM4) had a marginal effect on φ_exp_. In contrast, CIN85-BBB mixed with SUVs and either SLP65-R247A, SLP65_1-330_ or SLP65-3xPRM4 resulted in φ_exp_s of 1.1 ± 0.4 μM, 0.6 ± 0.3 μM and 1.0 ± 0.2 μM, respectively (Fig. S7*D-F*), all being lower by factors between 4 – 11 (φ ratios in Fig. 1*C*) compared to the φ_exp_ values measured with CIN85-ABC. Surprisingly, the CIN85-BBB construct, but not the SLP65-3xPRM4 construct, did promote tripartite phase separation, despite the fact, that SLP65 contains three strong binding modules, too, and both constructs having a lower K_D_ than the wildtype constructs (SLP65_1-330_, CIN85-ABC) (Table S1). This puts into evidence that K_D_s are not predictive for critical concentrations of phase separation and poses the question why triplication of PRM4 does not have the same phase separation promotion effect as introduction of the tripe SH3B construct.

### Replacement of SH3C by SH3B promotes phase separation properties of CIN85-BBB

As described above, the SLP65 mutants did not significantly shift the critical concentration in the tripartite system, but the CIN85 mutants did. We set out to understand these properties by simulations with the program LASSI taking into account the ranking of the individual K_D_s. Since we could not implement vesicles in LASSI, we investigated the SLP65-CIN85 system without SUVs.

Previous reports show that larger φ_exp_s are measured, when the vesicles, one of three essential components for the tripartite system, were omitted [14]. The mixtures of SLP65_1-330_ with CIN85-ABC resulted in φ_exp_ of 36 μM (Fig. 2*D*, Fig. S8*A*), which is, as expected, larger than the φ_exp_ of 6.7 μM including SUVs (Fig. 2C). Next, we analyzed the effect of the constructs designed to enhance the SH3/PRM interaction. The mixture of SLP65-3xPRM4 with CIN85-ABC showed a larger φ_exp_ of 62 μM (Fig. 2*D*) demonstrating a weak effect of SLP65-PRM4. The weak-binding SLP65-R247A mutant did not phase separate with the CIN85-ABC construct up to equimolar concentrations of 280 μM (Fig. S9), which confirms the relevance of PRM4 for CIN85 binding and corroborates previous findings regarding its role in CIN85 binding and Ca2+ signaling [cp. Fig. 3A in Oellerich et al. [27]]. While the SLP65-R247A construct did not phase separate with CIN85-ABC, it did so when mixed with CIN85-BBB, resulting in a φ_exp_ of 12 μM (Fig. 2*D*, Fig. S8*E*). SLP65_1-330_ with CIN85-BBB resulted in φ_exp_ of 4 μM (Fig. 2*D*, Fig. S8*F*) reproducing the previous finding that CIN85-BBB compared to CIN85 ABC promoted phase separation strongly (φ ratios “ABC / BBB” in Fig. 2*C*). Next, SLP65-3xPRM4 with CIN85-BBB resulted in φ_exp_ of 4 μM (Fig. 2*D*, Fig. S8*G*) reproducing the modest effect of triplication of PRM4 as seen before in the presence of SUVs. In conclusion, the tripartite system yielded φ_exp_ values ranging from 0.6 μM to 6.7 μM (Fig. 2C), while the φ_exp_ of the two protein system ranged from 2.9 μM to above 260 μM (Fig. 2*D*). Replacing CIN85-ABC by CIN85-BBB induces the most dramatic reduction in critical concentration of phase separation, yet the effects are quantitatively larger without SUVs than with (Fig. 2*C&D*). The replacement of SLP65_1-330_ by SLP65-3xPRM4 had almost no effect in the presence of SUVs and a quantitatively much smaller effect without SUVs than replacement of CIN85-ABC by CIN85-BBB.

**Fig. 3:**
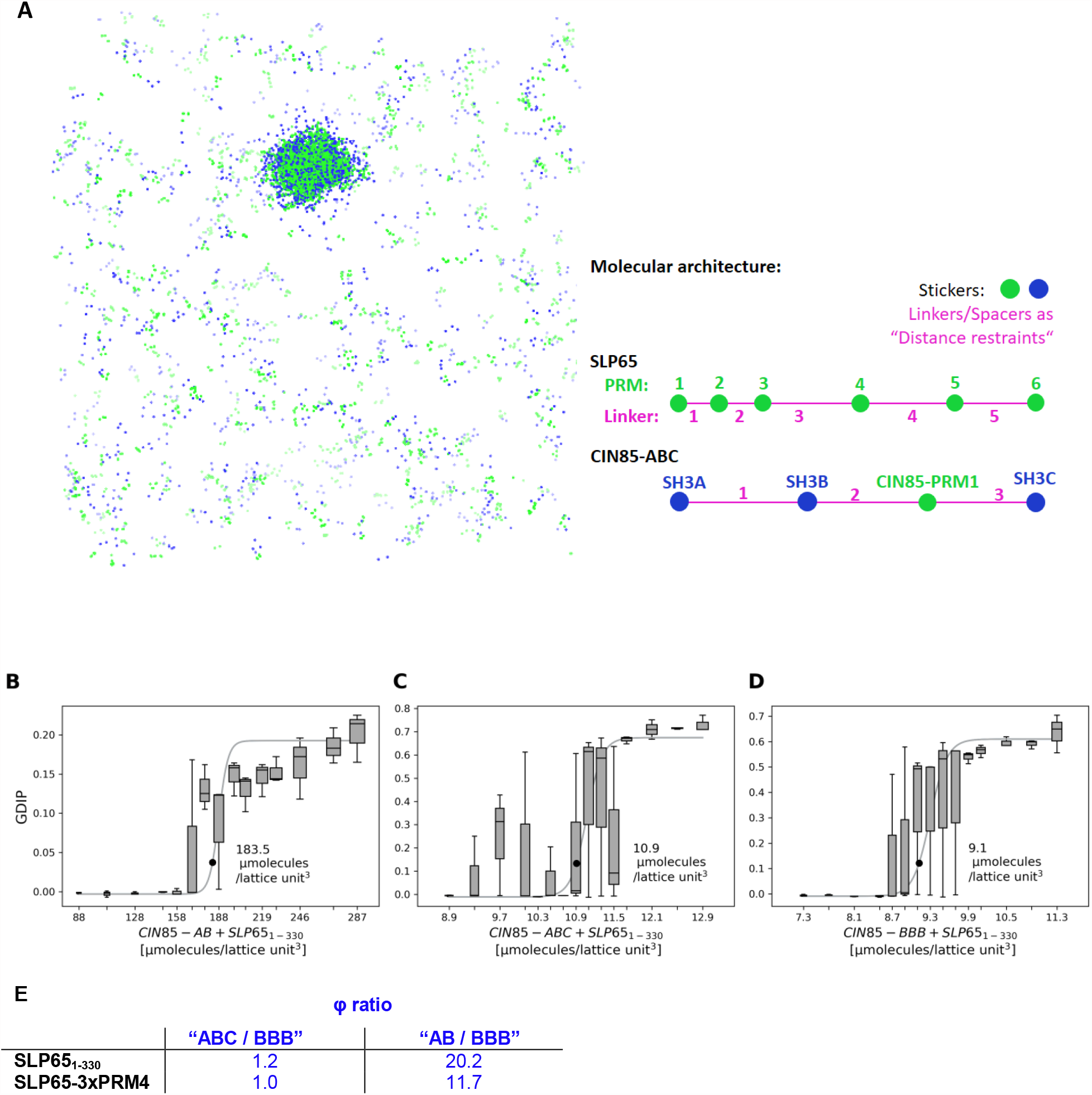
Analysis of CIN85’s and SLP65’s phase separation by simulations of the critical concentration φ_sim_. (A) LASSI simulations of the molecules’ PRM_m_-SH3_n_ system to visualize the molecular cluster of the SH3 stickers (blue) and PRM stickers (green) of 1000 *CIN85-BBB* molecules and 1000 *SLP65*_*1-330*_ molecules (2000 simulated molecules in total). In this representative simulation, the condensates are simulated at 9.1 μmolecules per lattice-unit^3^ and a spacer scaling = 1/6 (see main text). (B-D) *SLP65*_*1-330*_ was combined with either *CIN85-AB, CIN85-ABC* or *CIN85-BBB*. φ_sim_ was obtained by fitting the GDIP. The simulations with *CIN85-AB* (B) have a significantly different φ_sim_ than *CIN85-ABC* (C) and *CIN85-BBB* (D). E) φ ratios of the simulations (deduced from Fig. S13) with spacer scaling of 1/6).

Parallel to these studies, a new SH3 domain binding motif was identified in the linker between SH3B and SH3C, referred here as CIN85-PRM1 (D. Sieme et al., BioRxiv). We were interested, if CIN85-PRM1 could influence phase separation. The binary K_D_s of CIN85-PRM1 to SH3A, SH3B and SH3C were 720 ± 20 μM, 1140 ± 50 μM and 210 ± 10 μM as measured by NMR titration spectroscopy (Fig. 1*C*) (D. Sieme et al., BioRxiv). Since the SH3C domain is in close proximity to CIN85-PRM1, has the smallest K_D_, and occupies fully the PRM binding site of SH3C, CIN85-PRM1 is expected to prevent binding of SH3C to SLP65 PRMs. Indeed, probing CIN85-AB with SLP65_1-330_ resulted in a φ_exp_ of 18 μM (Fig. S8*C*, 36 μM for CIN85-ABC, Fig. 2*D*) and with SLP65-3xPRM4 in a φ_exp_ of 32 μM (Fig. S8*D)*, 62 μM for CIN85-ABC, Fig. 2*D*. The absence of SH3C and its linker containing CIN85-PRM1 rather promotes phase separation at concentrations lower by a factor of 0.5 (Fig. 2*D*). In conclusion, in CIN85-ABC, SH3C appears deactivated, such that it behaves very similar to CIN85-AB. CIN85-BBB, however, engages all three SH3B domains in interactions with PRMs of SLP65 since SH3B is not deactivated by CIN85-PRM1.

### Modeling of SLP65-CIN85 phase separation indicates that CIN85-ABC is effectively bivalent

We further elucidated the phase separation behavior of the SLP65 and CIN85 constructs introduced above by computational modelling with the program LASSI in order to relate the individual K_D_s to the phase separation behavior [25]. The lattice-based Monte-Carlo simulation uses stickers for which pairwise affinities can be defined and spacers which are neutral regarding binding (Fig. 3*A*). The SH3 domains of CIN85 and the PRMs of SLP65 constitute the stickers, whose interactions are defined by a sticker-sticker energy matrix parameterized by the experimentally determined ΔG values, derived from the K_D_ values (Fig. 1*C*, Fig. S10). Spacers represent the intrinsically disordered regions between the SH3 domains in CIN85 and between the PRMs in SLP65.

For simulations of CIN85, we included in addition to the CIN85-ABC and CIN85-BBB construct also the CIN85-AB construct. This is due to the fact that LASSI modelling could not reproduce the experimentally found 100% occupancy of the intramolecular interaction between SH3C and CIN85-PRM1 (Ref. D. Sieme, BioRxiv). Thus, CIN85-ABC with its inactive SH3C domain is therefore best represented by CIN85-AB.

Hereinafter we represent constructs for simulation in *italic* while constructs in experiments are shown in normal font. As a starting point, the spacer lengths were calculated in the following way. The PRMs are 6 amino acids long and represent one sticker lattice point. Therefore, the number of amino acids for a spacer was divided by 6 (scaling factor 1/6) to obtain the number of lattice points for a spacer (e.g. a 60 amino acid long linker is scaled to a spacer length of 10). In order to evaluate the impact of the spacer length we varied the scaling factors between 1/2 to 1/10 (Table S2). For illustration, the molecular cluster of the simulation of *CIN85-BBB* with *SLP65* is visualized (Fig. 3*A*). The simulated critical concentration (φ_sim_) for phase separation was determined by fitting the global density inhomogeneity parameter (GDIP), which reports about the dispersion of the molecules. A GDIP value above 0.025 indicates phase separation [25]. The φ_sim_ values of *SLP65* with *CIN85-AB, -ABC*, and *–BBB* are indicated as black point in Fig. 3*B-D*, respectively. In some simulations, a phase transition occurred at very high concentrations, where the GDIP was not suitable for our analysis any more. In this case, the percolation value, defined as the fraction of polymers of the single largest cluster [25], was used to calculate the critical concentration. In our simulations, the GDIP scales with the percolation value except for the simulations of *SLP65*_*1-330*_ with *SH3-AB* (Fig. S11).

The simulations of *CIN85-BBB* with either *SLP65* or *SLP65-3xPRM4* resulted in φ_sim_s of 9.1 and 9.4 μmolecules per lattice-unit^3^, respectively (Fig. S13, scaling 1/6), which were similar to the φ_sim_ values of 10.9 and 9.5 μmolecules per lattice-unit^3^ for *CIN85-ABC* and did therefore not match with the experimental observations in which CIN85-BBB lead to lower critical concentration than CIN85-ABC or CIN85-AB (Fig. 2D). The experiment and the simulation of CIN85-ABC did not agree (Figure 4, red circle). This non-matching behavior is expected when in the experiment SH3C does not participate in the interaction with PRMs of SLP65. In line with the deactivation of SH3C by CIN85-PRM1, the experimental φ_exp_-ratios of CIN85-AB *vs* CIN85-BBB (4.4 and 8.8, Fig. 2D, *colored blue*) matched with the simulated ones (20.2 and 11.7, Fig. 3*E*) within factors of 4.6 and 1.3, respectively. Thus, CIN85-ABC can be modelled faithfully by *CIN85-AB*, i.e. omitting the SH3C interactions with the SLP65 PRMs due to the binding of CIN85-PRM1 to SH3C.

**Fig. 4:**
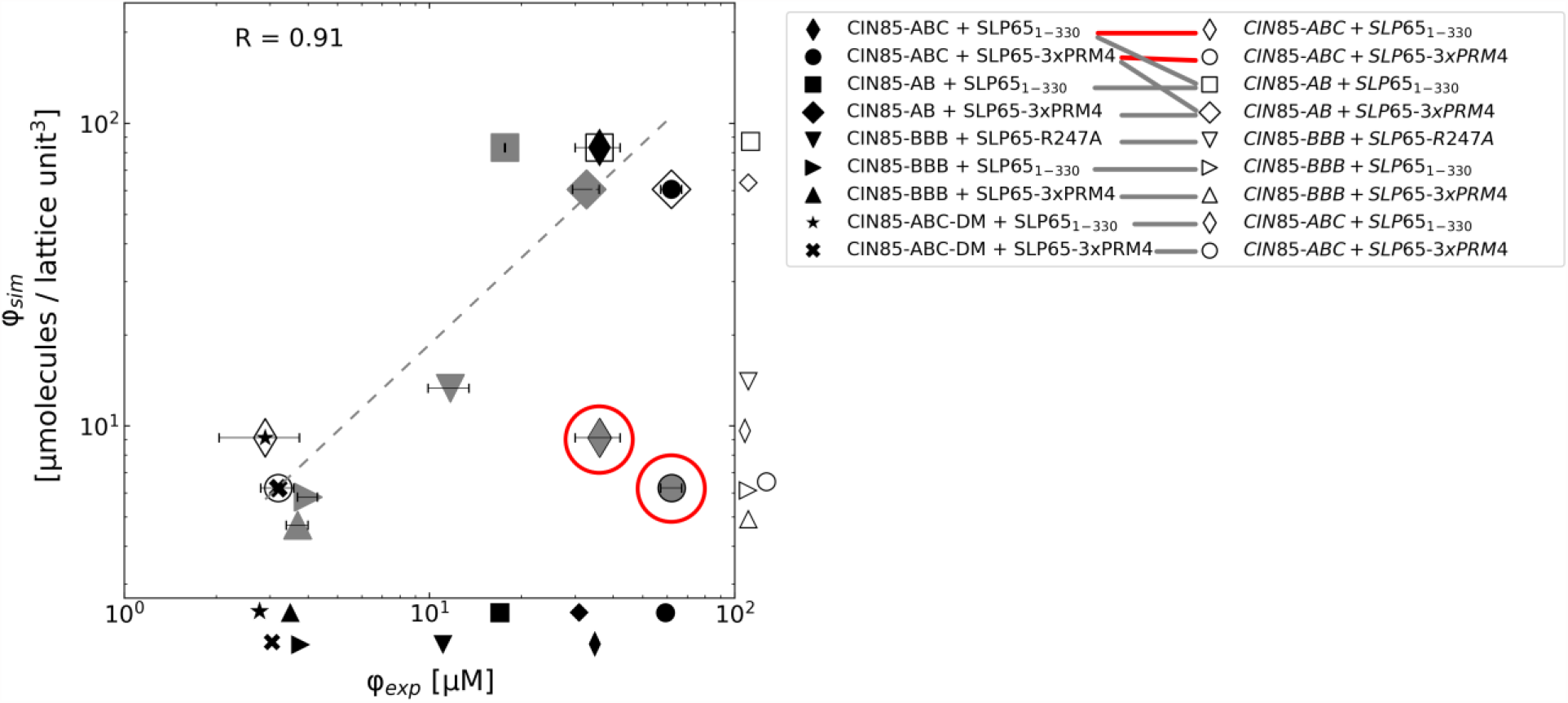
Correlation plots of φ_exp_-*vs* φ_sim_-values. *CIN85-AB* and *CIN85-BBB* correlate with CIN85-AB and CIN85-BBB, respectively. *CIN85-ABC* does not correlate with CIN85-ABC (circled red), but *CIN85-ABC* correlates with the CIN85-PRM1-inactivated CIN85-ABC-DM. The Pearson correlation coefficient is calculated from nine pairs of φ_exp_-*vs* φ_sim_-values, excluding the correlations of CIN85-ABC + SLP65_1-330_/SLP65-3xPRM4 with *CIN85-ABC + SLP65*_*1-330*_/*SLP65-3xPRM4* (circled red) φ-values of simulations with the spacer scaling of 1/4 are shown.

### The intramolecular interaction of CIN85-PRM1 impairs SH3C’s contribution to phase separation

The parameterized model showed a correlation of the simulated and experimental φ-values (Figure 4, white and black symbols, respectively) of CIN85-AB and CIN85-BBB (squares and triangles, respectively). We further investigated the contribution of CIN85-PRM1 to phase separation. We engineered a construct with inactivated CIN85-PRM1 in order to test its effect experimentally. CIN85-PRM1 was deactivated in the CIN85 double mutant (CIN85-ABC-DM) by the R227A and R229A mutations (D. Sieme, BioRxiv), i.e. it should be faithfully modelled by *CIN85-ABC*. The mixtures of SLP65_1-330_ with CIN85-ABC-DM (Fig. 4, black star) resulted in φ_exp_ of 2.9 μM (Fig. 2D; Fig. S8*H*) and correlates with the simulated *CIN85-ABC* (Fig. 4, white diamond). In contrast, CIN85-ABC with SLP65_1-330_ has a larger φ_exp_ of 36 μM and does not correlate with φ_sim_ (diamond in grey marked by the red circle), due to the CIN85 SH3C-PRM1 interaction not modelled in the LASSI simulations. φ_exp_ of CIN85-ABC-DM is indeed similar to the φ_exp_ of CIN85-BBB (Fig. 2*D*). This is recapitulated in the simulations: *CIN85-ABC* that ignores the intramolecular CIN85 SH3C-PRM1 interaction has a φ_sim_ of 10.9 similar to 9.1 of *CIN85-BBB* (Fig. S13). Similar to SLP65_1-330_ with CIN85-ABC-DM (2.9 μM, Fig. S8*H*), the mixture of SLP65-3xPRM4 with CIN85-ABC-DM has a φ_exp_ of 3.2 μM (Fig. S8*I*), which also agrees with the LASSI simulation: 9.5 for *CIN85-ABC* with *SLP65-3xPRM4* and 9.4 for *CIN85-BBB* with *SLP65-3xPRM4* (Fig. 4). The correlations were consistent with varying linker length scaling (Fig. S13 and S14). In summary, the mixtures of the CIN85-ABC-DM construct phase separated at critical concentrations similar to the mixtures of the CIN85-BBB construct (Fig. 4, bottom left part of the diagram), while the bivalent CIN85-AB construct and the CIN85-ABC construct have more similar phase separation properties (Fig. 4, upper right part) due to the deactivation of SH3C by CIN85-PRM1. CIN85-ABC-DM and the CIN85-BBB constructs are trivalent, whereas the CIN85-AB is bivalent and CIN85-ABC acts like a bivalent molecule. Supported by the LASSI model, the experiments demonstrate that valency is a major determinant for CIN85’s phase separation. This finding is in line with earlier studies, which revealed that phase separation of multivalent proteins is primarily governed by valency [24, 31-33], while further determinants driving phase separations such as affinity [34] and linker length [35, 36] are secondary.

## Conclusions

Computer simulations implementing interactions by the sticker-spacer concept have been used to predict sequence patterns mediating the phase separation of IDPs [32, 35]. We applied the LASSI model to the natural SH3-PRM system of SLP65 and CIN85, and described the phase separation process emerging from natural, promiscuous PRM-SH3 interactions. The phase separation concentrations could be predicted from the individual dissociation constants of the individual components faithfully and variations of composition were described correctly by the LASSI simulations. LASSI was further used as a heuristic tool to detect previously undetected interactions: while analyzing the affinity-dependent SLP65/CIN85 phase separation, we noticed a discrepancy between the simulated and measured critical concentrations of the CIN85-ABC constructs (Fig. 4, circled red). This was in line with the identification of bivalency in CIN85-ABC, where SH3C was deactivated by intramolecular binding to CIN85’s PRM1 (Sieme, BioRxiv). These findings highlight the impact of valency on the critical concentration. Moreover, the simulations comprehensibly describe configurations of a phase separated, large molecular cluster at a protein domain level resolution.

## Materials and Methods

### Reagents

Peptide synthesis and vesicle preparation, as well as protein production can be found in the Material and methods

### Biophysical interaction studies

NMR and ITC titration experiments are described in the supplementary information.

### Fluorescence microscopy assay and image analysis

Sample preparation was performed as described previously [14]. Briefly, the protein constructs were mixed with or without SUVs in 1.5 ml reaction tubes (Eppendorf). The total volume of each mixture was 25 μl. For a robust assay, the components were added in the distinct order, starting with the addition of buffer (20 mM HEPES pH 7.2, 100 ml NaCl). Then, the atto430LS-labeled SLP65 construct, the unlabeled SLP65 construct, and the CIN85 construct were added sequentially. For mixtures including SUVs, a SUV stock solution with 5 mM lipid concentration was pipetted to the buffer first, and then the protein constructs were pipetted. The mixtures were transferred into the wells of a microscopy slide (uncoated μ-Slide 8 Well, ibidi GmbH, Gräfelfing). Before imaging, phase separated droplets were allowed to settled on the surface during an incubation time of 45 – 75 minutes. The phase separation of mixtures with SLP65, CIN85 and SUVs were imaged with a Zeiss LSM780 confocal microscope equipped with an MBS 458 beam splitter and a Plan-Apochromat 40x/1.4 Oil DIC M27 objective. To image the settled dense phase right above the bottom of the well, a 4-8 μm z-stack with a step size of 1 μm was acquired. Atto 430LS-labeled SLP65 was excited by an argon laser at 458 nm. Emission was detected in the 520-600 nm range. Laser settings and representative acquisition parameters were chosen depending on the fluorophore concentration. For quantification and to avoid intensity saturation of the image, not more than 10 μM atto430LS-labeled SLP65 was used. In parallel, bright field images were recorded (detector gain BF: 400). Images of the surface (z-position at _∼_0 μm) were selected and analyzed with the ImageJ software (NIH, Bethesda, MD, USA). The dispersion index σ^2^ / μ was calculated from 16-bit images (n = 15), where σ is the variance and μ the mean of the pixel intensities. The critical concentration φ_exp_ for phase separation was obtained by fitting the dispersion index to the logistic function 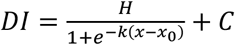 where H, C and k are scaling parameters of which H and C depend on the absolute signal intensity, x0 is a location parameter, and φ_exp_ is defined as the maximum of the 2nd derivative of the fit.

### Computational model

In the applied LASSI model [25], the definitions of SLP65’s and CIN85’s molecular architecture are described in the main text (Fig. 3) and illustrated in the Table S2 (supplementary information). The simulations included in total 2000 molecules (1000 for each SLP65 and CIN85) in order to avoid finite-size effects [25]. Rotational, local, co-local, shake, translation, small cluster translation, cluster translation, pivot and double pivot moves were applied at frequencies of 0.25, 0.13, 0.25, 0.04, 0.04, 0.04, 0.04, 0.13, 0.10 per step considering the suggestions for the settings from the authors [25]. The anisotropic interaction energy terms were experimentally parametrized by the Gibbs free energy ΔG (Fig. S10) in terms given by ΔG = -RT ln(1/K_D_), with the dissociation constant K_D_, the temperature T = 310 K and the gas constant R = 1.987 cal mol^-1^ K^-1^. The isotropic interaction energy terms were set to zero to allow for simulations with short linker lengths as suggested by the authors of LASSI (personal correspondence). 2*10^9^ Monte-Carlo steps were run in order to approach apparent convergence of the simulation. To simulate the critical concentration, a set of simulations were performed, in which the concentration was varied by reducing the box size. In the rectangular cuboid box, the simulated concentration is given in units of μmolecules per lattice-unit^3^, which represents 10^−6^ molecules per lattice-unit^3^, with 2000 molecules and a box size length between 85 to 900 lattice units. The radial density distribution (provided by the software) was normalized with the a priori radial density distribution (simulations with zero energy interaction terms). The GDIP was calculated from the normalized radial density distribution function for each simulation. The critical concentration was obtained by fitting the GDIP of three simulations (n = 3). Bootstrap errors were calculated for the simulated and experimentally determined critical concentration with 5000 repetition. For each repetition, 15 dispersion index values or 3 GDIP values of the sample data were randomly chosen per concentration point, and fitted to obtain the critical concentration. The standard deviation was calculated from the list of the critical concentrations.

## Author Contributions

J.M. S.B., and C.G. designed research; J.M., D.S. and L.E.W. performed research; J.M., D.S. and F.D. analyzed data; J.W. and C.G. secured the funding for the research; J.M., S.B., D.S. and C.G. wrote the paper; S.B. contributed reagents.

## Acknowledgements

We thank Claudia Schwiegk and Kerstin Overkamp for excellent technical help and peptide synthesis. We thank Ángel Pérez-Lara for assistance with ITC experiments and the Live-cell imaging facility at the MPI for Multidisciplinary Sciences for access to the microscopes. This project was supported by the DFG (Deutsche Forschungsgemeinschaft) in the framework of the SFB860.

## Notes

### Competing Interest Statement

The authors have declared no competing interest.

