## Supplementary Information for "Quantitative description of the phase separation behavior of the multivalent SLP65-CIN85 complex"

Christian Griesinger

Mail:

### Methods and Materials

##### Peptide synthesis

The SLP65 and CIN85 peptides were synthesized by solid-phase peptide synthesis with an acetylated N-terminus and amidated C- terminus. Peptides were purified on a RP18 column in $H_2O$ with 0.1 % TFA and eluted in acetonitrile with 0.1 The lyophilized powder was diluted in a 100 mM NaCl, 20 mM HEPES buffer and the pH was adjusted to obtain a stock solution at pH 7.2.

##### Vesicle preparation

The preparation of SUVs was adapted from the previously described procedure [1]. Lipids were weighed in separately in glass vials (Duran) and completely resuspended in chloroform to obtain DOPC, DOPE and DOPS stock solutions. The stock solutions were mixed in the ratio 65:25:10 (DOPC: DOPE: DOPS *%w/w*) to obtain a mixture with a total volume of 1 ml and a final total lipid concentration of 5 mM. A lipid film was prepared by drying the lipid composition under a nitrogen stream, followed by lyophilization. The lipid film was resuspended in 20 mM HEPES (pH 7.2) buffer with 100 mM NaCl, rigorously shaken for complete resuspension and transferred to a 15 ml plastic tube (Falcon). Vesicles were formed by high power tip sonication on ice for 2 x 20 min, including a break to decrease the temperature of the sonicated suspension (Sonoplus HD 3100). For tip sonication a 60% amplitude power was set and pulsation switched in a 2-3 sec on and 2 sec off. After sonication, the suspension was diluted 1:10, in order to measure the vesicle hydrodynamic radius distribution by dynamic light scattering (DLS). The suspension was checked for clarity and then further extruded by transferring the suspension (Mini Extruder, Avanti Polar Lipids) 25 times through a 50 nm polycarbonate membrane filter (Avestin) to achieve a more homogenous vesicle size distribution. To check the extrusion procedure, the SUV hydrodynamic radius distribution of a 1:10 dilution was measured by DLS. The 5 mM-lipid SUV preparation was diluted 1:2 to a obtain a SUV stock suspension with 2.5 mM lipids.

##### Recombinant protein production

Plasmids encoding SLP65-3xPRM4 and CIN85-BBB were purchased (Invitrogen) and the coding fragments were cloned into expression vectors. The SLP65-3xPRM4 fragment was inserted into a pET16b/TEV vector via BamH1/NdeI restriction sites. The $SLP65_{1\text{-}330}$-R247A (in the following referred to as SLP65-R247A) plasmid was generated by site-directed PCR mutagenesis on the $SLP65_{1\text{-}330}$ plasmid [1] using the forward primer $^{5'}$CAC CAT CCC CGT TGC CAG CGG CCG GGA AAA AAC CAA C$^{3'}$ and the reverse primer $^{5'}$GTT GGT TTT TTC CCG GCC GCT GGC AAC GGG GAT GGT G$^{3'}$. For fluorophore labeling, a C-terminal cysteine was introduced by site-directed mutagenesis using the primers $^{5'}$CCT CTG CCG AGC TTT AGC AGC TGC TAA AGG ATC CGG CTG CTA AC$^{3'}$ (forward) and $^{5'}$GTT AGC AGC CGG ATC CTT TAG CAG CTG CTA AAG CTC GGC AGA GG$^{3'}$ (reverse). The coding sequence for CIN85-BBB was cloned via

BamH1/HindIII restriction sites into a modified pET28a vector (Novagen) encoding for an N-terminal Z2 fusion domain with TEV cleavage site and a 7x histidine tag. All $SLP65_{1-330}$ and CIN85-ABC protein constructs were expressed in *E. coli* and purified by metal affinity chromatography as described previously [1].

The sequences of SH3A and SH3B were each cloned into a pGEX4-T vector, which encodes a glutathione-S-transferase (GST) fusion protein. Each fusion protein GST-SH3A and GST-SH3B was expressed in *Escherichia coli* BL21 (DE3) (New England Biolabs) and purified by affinity chromatography (resin: Pierce™ Glutathione Agarose). Elution fractions were collected and dialyzed at 23 °C against 20 mM HEPES (pH 7.2), 150 mM NaCl, 1 mM DTT. Thrombin was added during dialysis to cleave off the GST fusion tag. After cleavage, thrombin and the cleaved GST tag were removed by size exclusion chromatography (Superdex 75 16/60 GL, GE Healthcare). Fractions were pooled and dialyzed for >12 h at 4° C against 20 mM HEPES (pH 7.2), 100 mM NaCl buffer, 1 mM DTT and 0.5 mM PMSF. The protein samples were concentrated by centrifugation (Vivascience concentrators, Sartorius) and stored at – 80 °C until further usage.

The coding sequence of SH3C was cloned into a modified pET16b vector including a TEV-protease cleavage site and a 7 x histidine tag for affinity purification. The protein was expressed in *Escherichia coli* BL21 (DE3) (New England Biolabs) and purified by Ni-NTA affinity chromatography. The SH3C protein was dialyzed for > 12 h at 23 °C against 20 mM Tris/HCl (pH 8.0), 200 mM NaCl, 0.5 mM EDTA, 1 mM DTT, 0.5 mM PMSF buffer including TEV-protease (TEV-Protease:SH3C 1:50) to cleave off the 7 x histidine tag. To remove the cleaved 7 x histidine tag, a reverse affinity purification (Ni-NTA Protino resin, Macherey-Nagel) was done. Then, protein was dialyzed against 20 mM HEPES (pH 7.2), 100 mM NaCl, 1mM DTT, 0.5 mM PMSF and afterwards purified by size exclusion chromatography in the same buffer.

For fluorescence labeling, atto 430LS dye (Atto-Tec) was conjugated by a maleimide thiol reaction (according to the manufacturer´s protocol). $SLP65_{1-330}$, SLP65-R247A and SLP65-3xPRM were atto 430LS-labeled at residues C271, at the introduced mutation S236C and at the C-terminal added C331, respectively. Each reaction product was purified by size exclusion chromatography (Superdex 75 10/300 GL, GE Healthcare), respectively.

Protein concentrations were determined by UV-Vis spectroscopy using the molar extinction coefficient at 280 nm. For atto430LS labeled proteins, the absorption of the dye was subtracted using the absorbance ratio of 280 nm / 430 nm.

**NMR experiments**

$^{15}N$-labeled protein constructs were expressed in M9 minimal medium [2] supplemented with $^{15}N$-$NH_4Cl$ and expressed as described above. Samples were supplemented with 0.5 mM DSS and 4% $D_2O$ for chemical shift referencing and locking, respectively.

The NMR triple resonance assignment experiments HNCACB (hncacbgp3d, [3, 4]) and CBCA(CO)NH (cbcaconhgp3d [4, 5] were acquired of both 1 mM $^{13}C$,$^{15}N$-labeled SH3A domain and 0.5 mM $^{13}C$,$^{15}N$-labeled SH3C domain at 700 MHz and 800 MHz spectrometers, respectively, in 20mM HEPES (pH 7.2), 50 mM NaCl buffer at 298 K. For SH3B assignment, the HNCA (hncagpwg3d [6-8]). NMR experiments were recorded in 20mM HEPES, pH 7.2, 50 mM NaCl buffer at 600 MHz at 298 K. In order to analyze $^{15}N$-HSQC titration experiments, a $^{15}N$-HSQC experiment at 30 °C was recorded, and the $^{1}H$ and $^{15}N$ assignment were transferred from 25 °C to 37 °C by following the chemical shift with increasing temperature. The NMR titrations were performed at 37 °C on 600 MHz, 700 MHz, 800 MHz or 900 MHz spectrometers equipped with a z-gradient cryoprobe. The receptor concentration of the CIN85's SH3 domains and the titration ratios (ligand:protein) are listed in the Table S3. Binding of PRM-derived peptides to $^{15}N$-labeled SH3 domains were monitored by $^{15}N$-HSQC NMR experiments. For acquisition of the sensitivity-enhanced $^{15}N$-HSQC experiment, an adapted version of the standard pulse program (hsqcetf3gpsi, [9-11]) was used, with 2048 complex points in the direct dimension, 256 complex points in the indirect dimension, 4-12 scans and a recycle delay in the range of 1.2 – 1.5 s. A water-flipback pulse for water suppression and waltz16 decoupling was applied. Spectra were processed in Topspin 4.06 (Bruker) applying a cosine-squared window function in both dimensions and if required a $0^{th}$ or $1^{st}$ order phase correction.

NMR spectra were analyzed by 1D chemical shift or 2D lineshape analysis [12]. For 1D chemical shift analysis the $^{1}H$ and $^{15}N$ chemical shift perturbations were combined according to the following formula: $^{1}H$-$^{15}N_{combined} = \sqrt{(( CS_N * 0.1)^2 + CS_H^2}$, where $CS_N$ is the chemical shift difference in the nitrogen dimension and $CS_H$ is the proton dimension. The $K_D$ of a titration was obtained by a least square fit of the equation: $CS = \frac{\Delta \mathrm{CSmax}}{2*[\mathrm{P}]_0} * ((K_D + [\mathrm{L}]_0 + [\mathrm{P}]_0) - \sqrt{(K_D + [\mathrm{L}]_0 + [\mathrm{P}]_0)^2 - 4 * [\mathrm{L}]_0 * [\mathrm{P}]_0})$ , where $\Delta CS$ is the chemical shift difference of the observed CS and the CS of the protein in the unbound state. $\Delta CS_{max}$ is the maximum chemical shift difference, $L_0$ is the total ligand concentration (which is titrated) and $P_0$ is the total protein concentration (constant).

**ITC measurements**

The interactions between single SH3 domains and single peptides and the interactions of SLP65 constructs to CIN85 constructs were measured by isothermal ITC on a VP-ITC MicroCalorimeter (MicroCal). Titrations were done at 37 °C with 60 s initial delay, a stirring speed of 307 rpm and a reference power of 10 µCal/second. The injection scheme is shown in Table S4. For the measurement of monovalent interactions, 500 µM peptide was titrated to 20 µM SH3 domain with a spacing time between the injection points of 400 s. For titration of multivalent designer constructs, the spacing time was increased to 600 s to reach equilibration after each injection. For smoothening the data, the filter period over which the data channel conversions are averaged was set to 2 s. Peptides and protein samples of the SLP65 and CIN85 constructs, dissolved in 20 mM HEPES (pH 7.2) buffer with 100 mM NaCl, were degassed before titration.

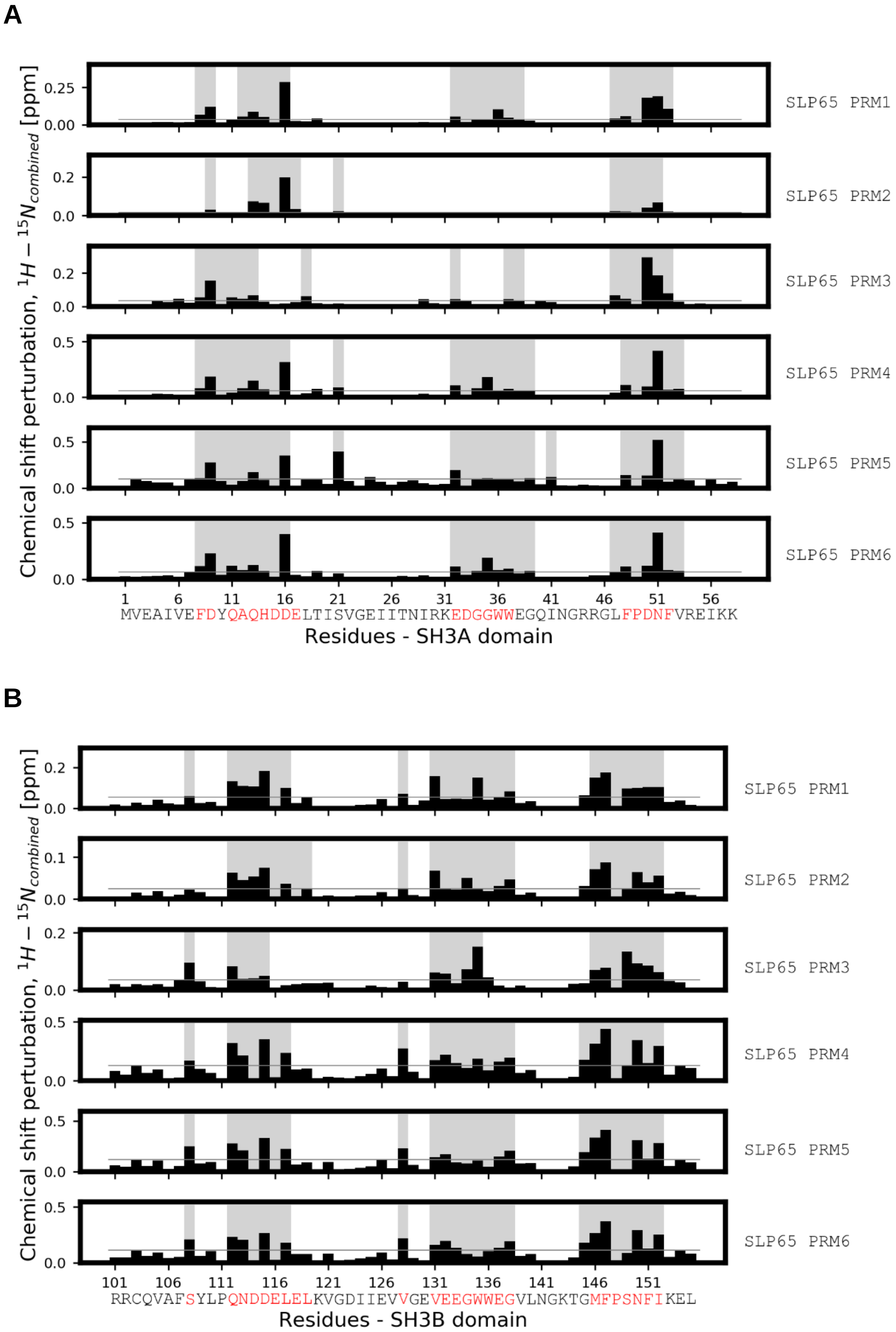
A
Chemical shift perturbation, $^1H - ^{15}N_{combined}$ [ppm]
SLP65 PRM1
SLP65 PRM2
SLP65 PRM3
SLP65 PRM4
SLP65 PRM5
SLP65 PRM6
1 6 11 16 21 26 31 36 41 46 51 56
MVEAIVEFDYQAQHDDELTISVGEIITNIRKEDGGWWEGQINGRRGLFPDNFVREIKK
Residues - SH3A domain
B
Chemical shift perturbation, $^1H - ^{15}N_{combined}$ [ppm]
SLP65 PRM1
SLP65 PRM2
SLP65 PRM3
SLP65 PRM4
SLP65 PRM5
SLP65 PRM6
101 106 111 116 121 126 131 136 141 146 151
RRCQVAFSYLPQNDDELELKVGDIIEVVGEVEEGWWEGVLNGKTGMFPSNFIKEL
Residues - SH3B domain

**C**

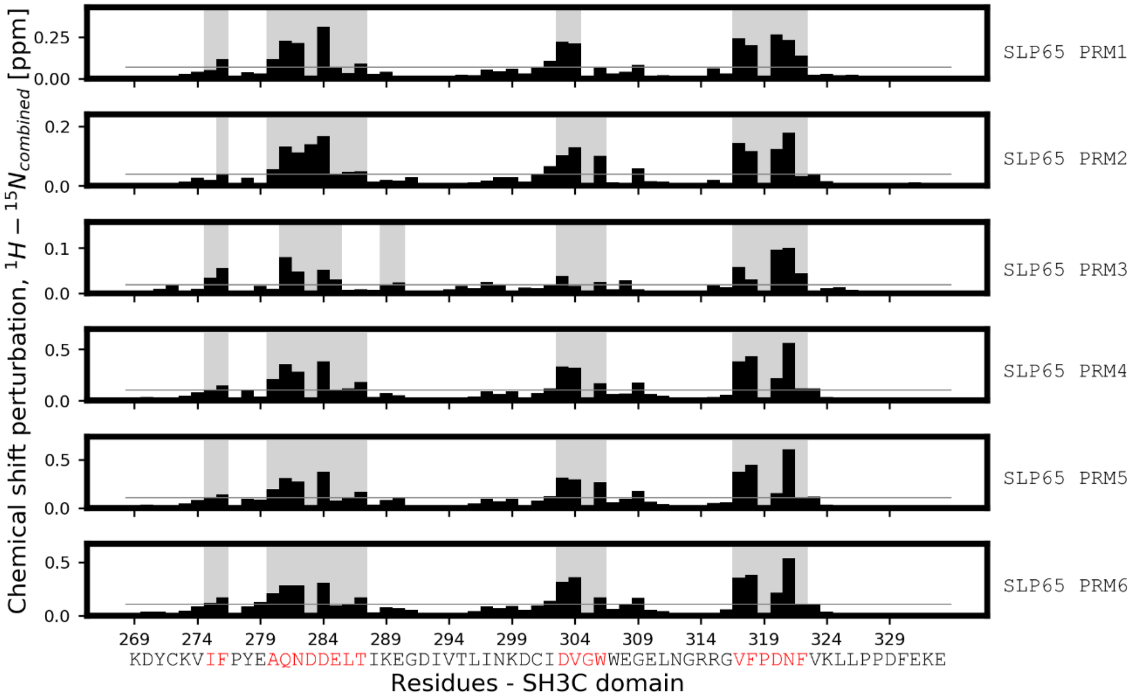


Fig. S1: Chemical shift perturbations of titrations of SLP65-PRM1-6 to the SH3A (A), SH3B (B) and SH3C domain (C). The average CSP of each titration is indicated as grey line. Residues and regions with large CSP are shaded grey and highlighted in the sequence (red). For SH3A, the RT-loop, N-src loop and β4-strand). For SH3B, the regions with large CSP were in particular the RT-loop, the N-src loop, β4-strand and the $3_{10}$-helix. For SH3C, the RT-loop, the N-src loop, β4-strand and the $3_{10}$-helix. Each SH3 domain has a similar CSP profile indicating a single, common binding site. The CSP are reported for the endpoint of the titration with a ligand/receptor ratio of 3.3, 4.7, 4.3, 10.7, 9.0 and 9.0 (A), 4.0, 10.0, 5.6, 10.0, 8.0, 4.7 (B) and 9.0, 12.8, 9.0, 9.0, 9.0 and 9.0 (C) for SLP65-PRM1, 2, 3, 4, 5, 6, respectively (Table S3).

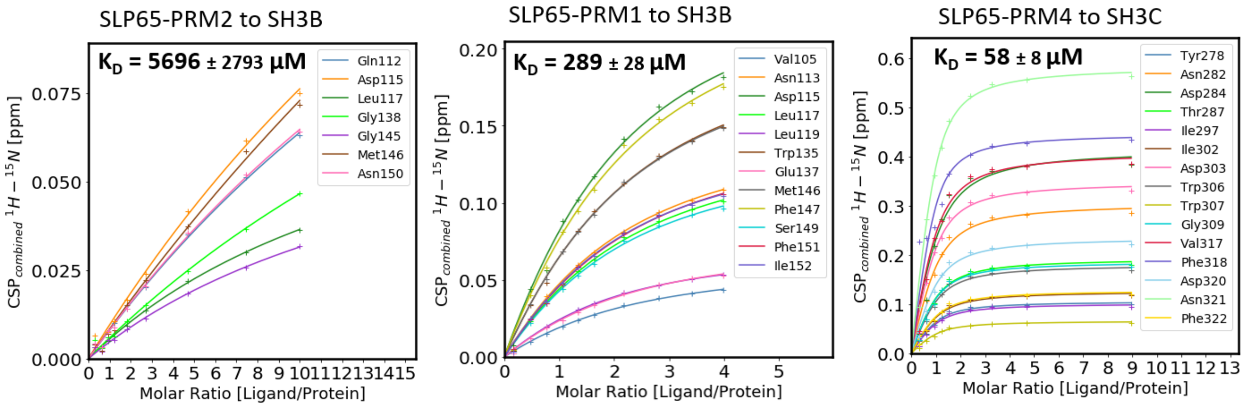


Fig. S2: Representative binding isotherms of the combined chemical shift analysis of a weak, medium and strong PRM-SH3 domain interaction. SLP65-PRM1-6 were titrated to either SH3A, B or SH3C. The combined chemical shift was fitted to obtain the dissociation constants of SH3 domain-PRM titrations. Representative binding isotherms of titrations of SLP65-PRM2 and SLP65-PRM1 to the SH3B domain and SLP65-PRM4 to the SH3C domain are shown.

A

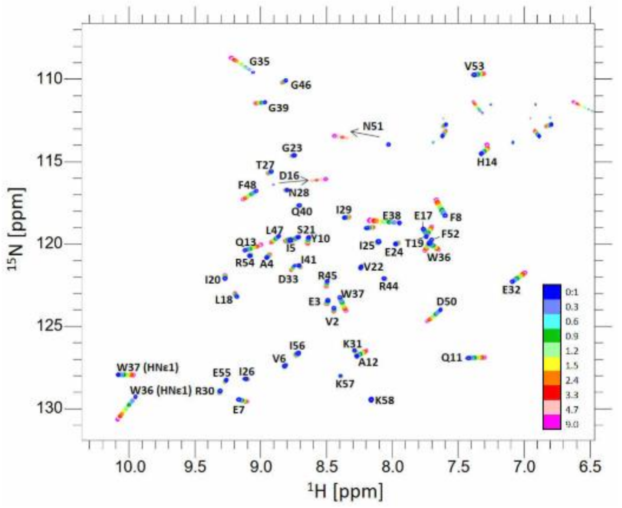


B

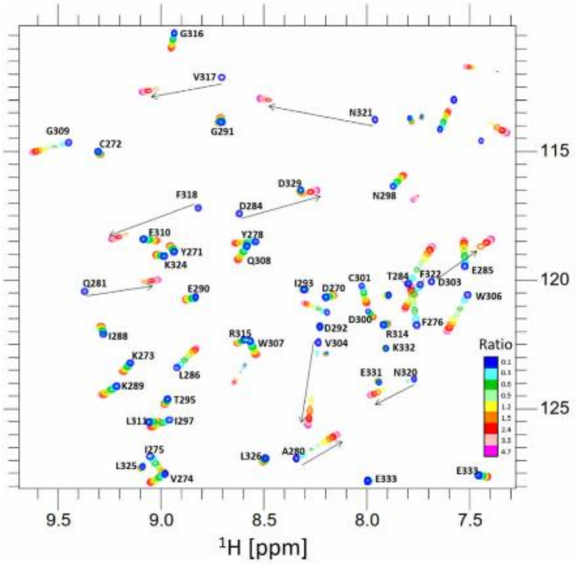


C

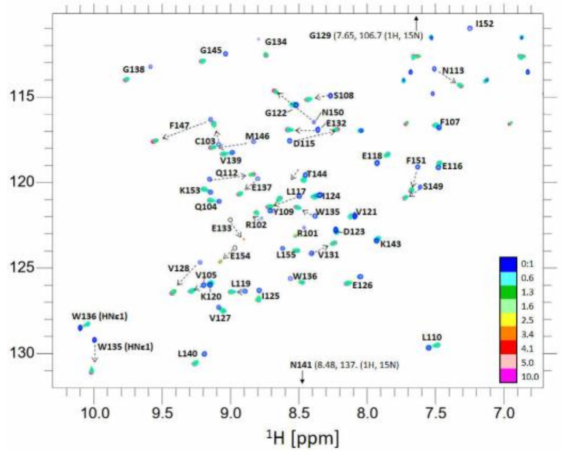


Fig. S3: Overlaid NMR spectra of the titrations of SLP65-PRM6 to the $^{15}$N-SH3A (A), SLP65-PRM4 to $^{15}$N-SH3C (B) and SLP65-PRM4 to $^{15}$N-SH3B (C). Representative spectra for different exchange regimes from the NMR titration experiments of the SLP65-peptide interaction with either $^{15}$N-SH3A, $^{15}$N-SH3B or $^{15}$N-SH3C domain are shown. The titrations display the majority of the peaks in fast (A), intermediate (B, marked with arrows) and slow (C) exchange regime. $^{15}$N-HSQC spectra were recorded at 800 MHz at 37°C and an initial SH3 domain concentration of 200 µM (0:1 ligand:protein ratio). Titration spectra are colored according to the ligand to protein ratio.

Titration: **PRM1 to SH3B** **PRM5 to SH3C** **PRM4 to SH3B**

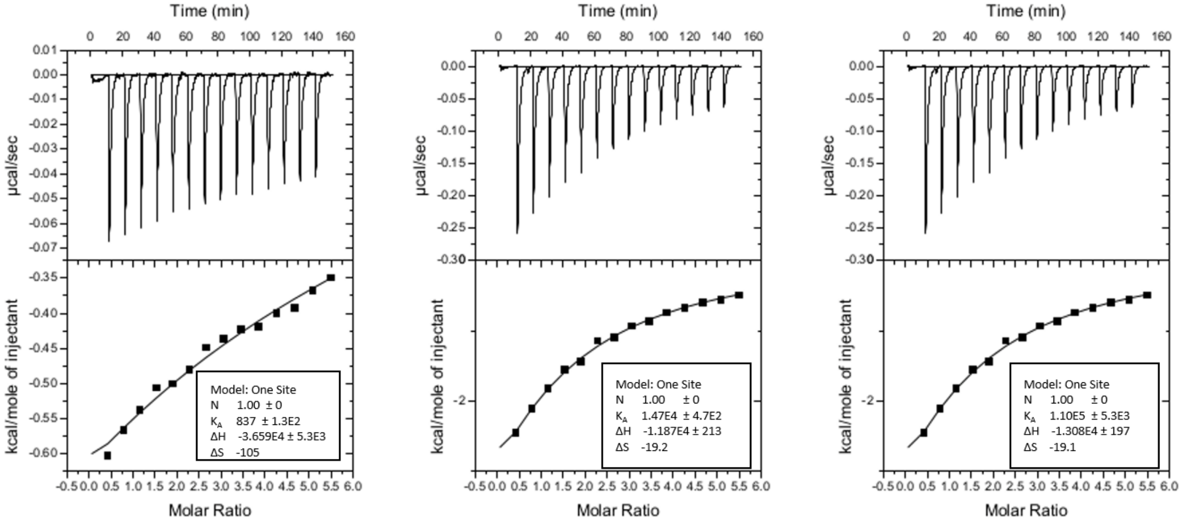


Titration: **PRM2 to SH3A** **PRM2 to SH3B** **PRM2 to SH3C**

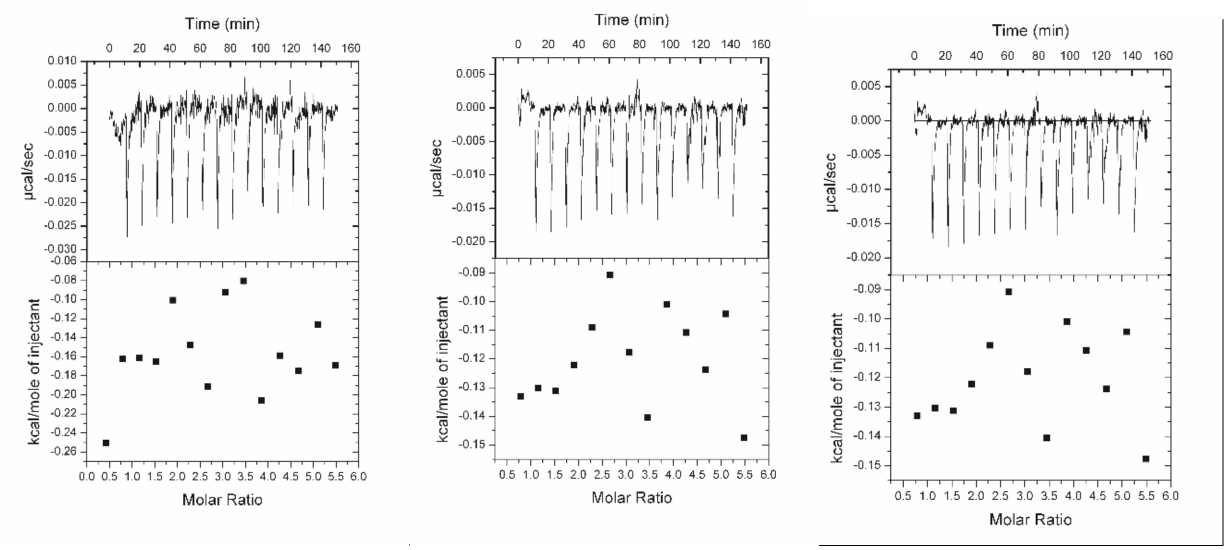


Fig. S4: Representative ITC binding isotherms of a weak, medium and strong PRM-SH3 domain interaction. The interactions of PRM1 to SH3B (top left), PRM5 to SH3C (top middle) and PRM4 to SH3B (top right) are shown. No heat evolution was measured for the titration of PRM2 to the SH3A, -B or -C domain (lower panel).

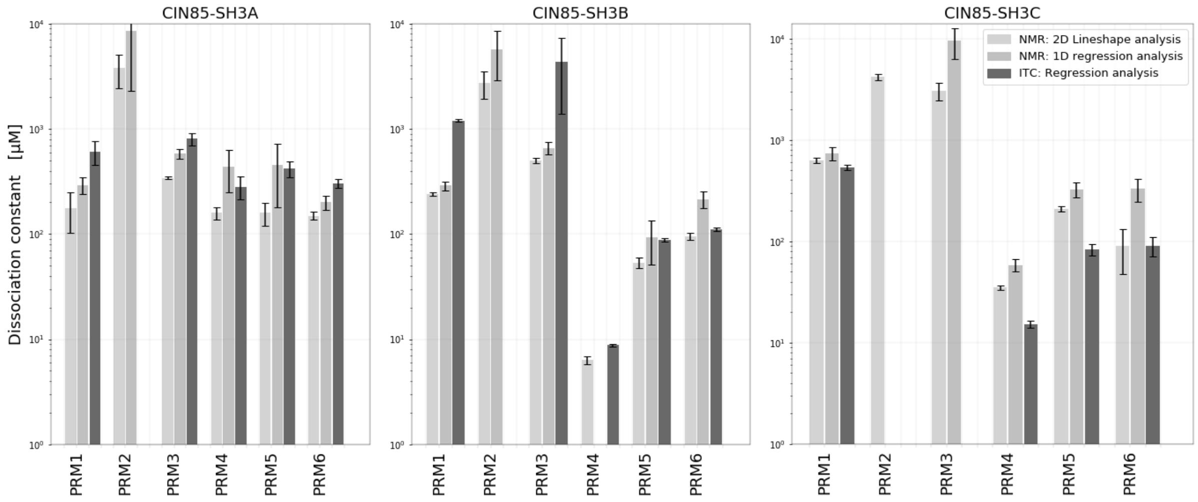


Fig. S5: Dissociation constants ($K_D$s) of the interactions of SLP65-peptides with any of the three CIN85-SH3 domains. Comparison of the $K_D$s measured by ITC and NMR. NMR data was both analyzed by fitting of the chemical shift perturbation (colored lightgrey, 1D regression analysis) and by 2D lineshape analysis (colored silver) using the program TITAN v1.5-3-g3566 [12]. The ITC analysis is colored dim grey.

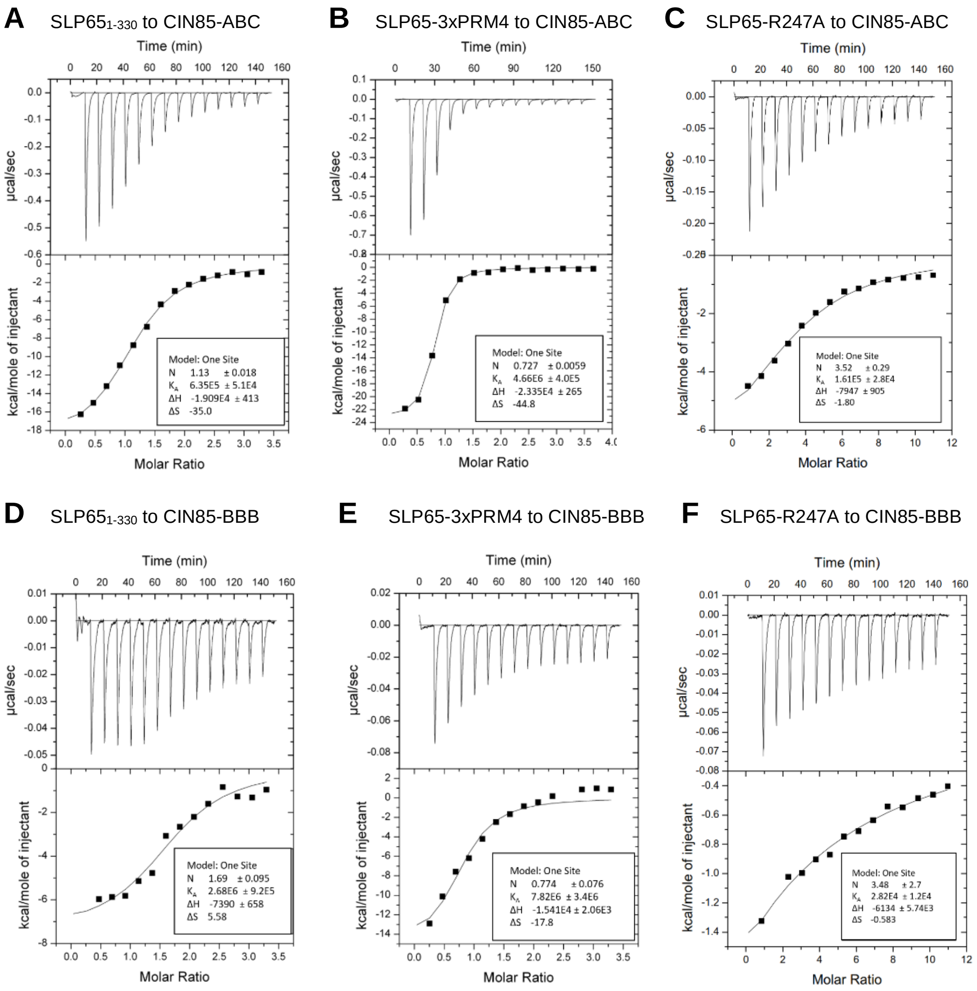


Fig. S6: Representative ITC binding isotherms of interactions of SLP65$_{1-330}$ (150 µM syringe concentration) to 10 µM CIN85-ABC (A), SLP65-3xPRM4 (170 µM syringe concentration) to 10 µM CIN85-ABC (B), SLP65-R247A (500 µM syringe concentration) to 10 µM CIN85-ABC (C), SLP65$_{1-330}$ (30 µM syringe concentration) to 2 µM CIN85-BBB (D), SLP65-3xPRM4 (15 µM syringe concentration) to 1 µM CIN85-BBB (E) and SLP65-R247A (150 µM syringe concentration) to 3 µM CIN85-BBB (F). Larger and smaller $K_D$s of the weak- and strong-binding versions, respectively, were confirmed.

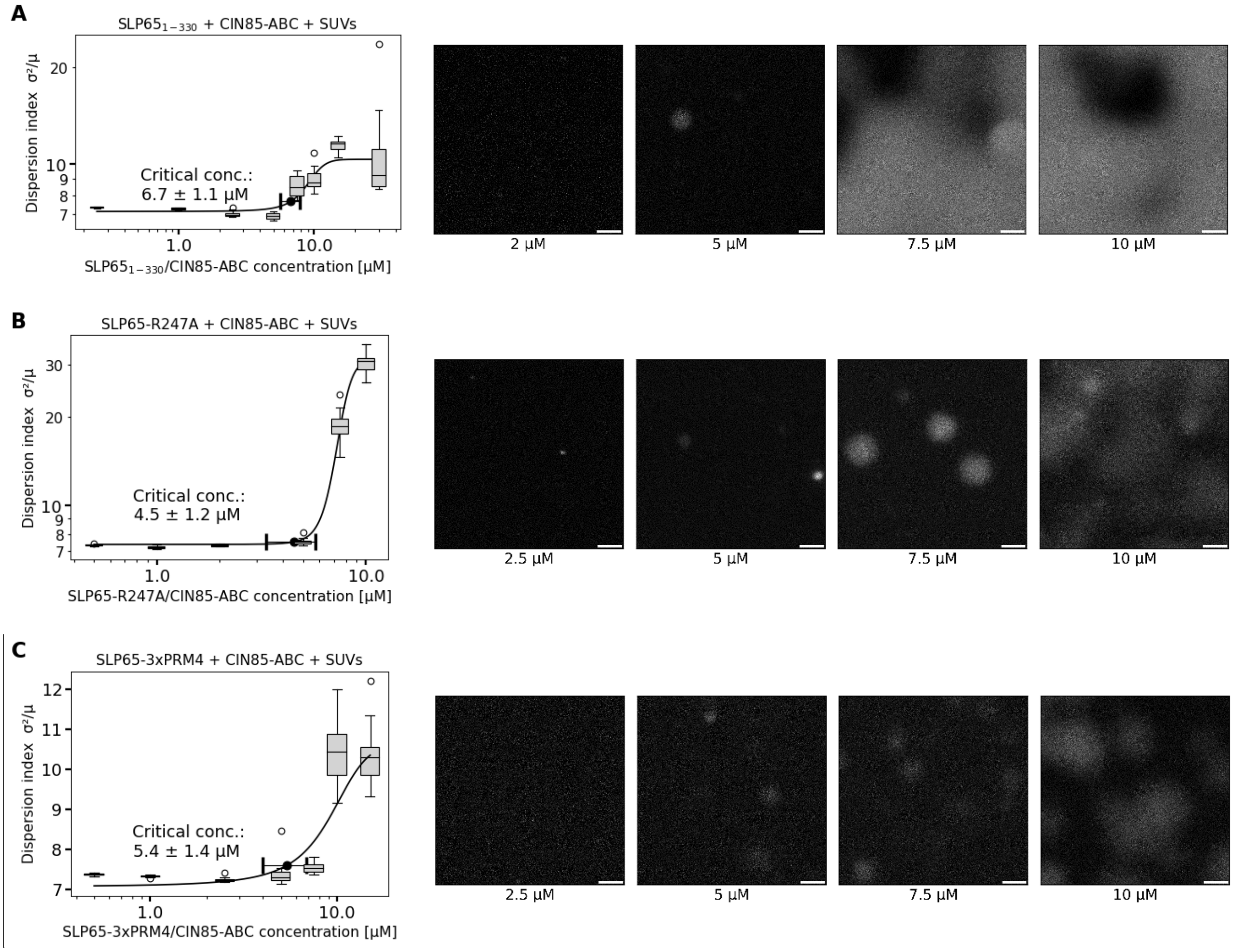


Fig. S7: Measurement of critical concentrations of SLP65-CIN85-SUV condensates by confocal fluorescence microscopy: Mixtures were prepared at equimolar SLP65/CIN85 concentrations. Different-affine SLP65 constructs were probed. A) Mixtures containing SLP65$_{1\text{-}330}$, CIN85-ABC and SUVs. B) Mixtures of SLP65-R247A, CIN85-ABC and SUVs. C) Mixture of SLP65-3xPRM4, CIN85-ABC and SUVs. The critical concentration with bootstrap SD errors (highlighted in green) were obtained from fitting the dispersion index (left). Representative images (right) of mixtures show no phase separation at low concentration and phase separation at high concentrations. Scale bar: 10 µm.

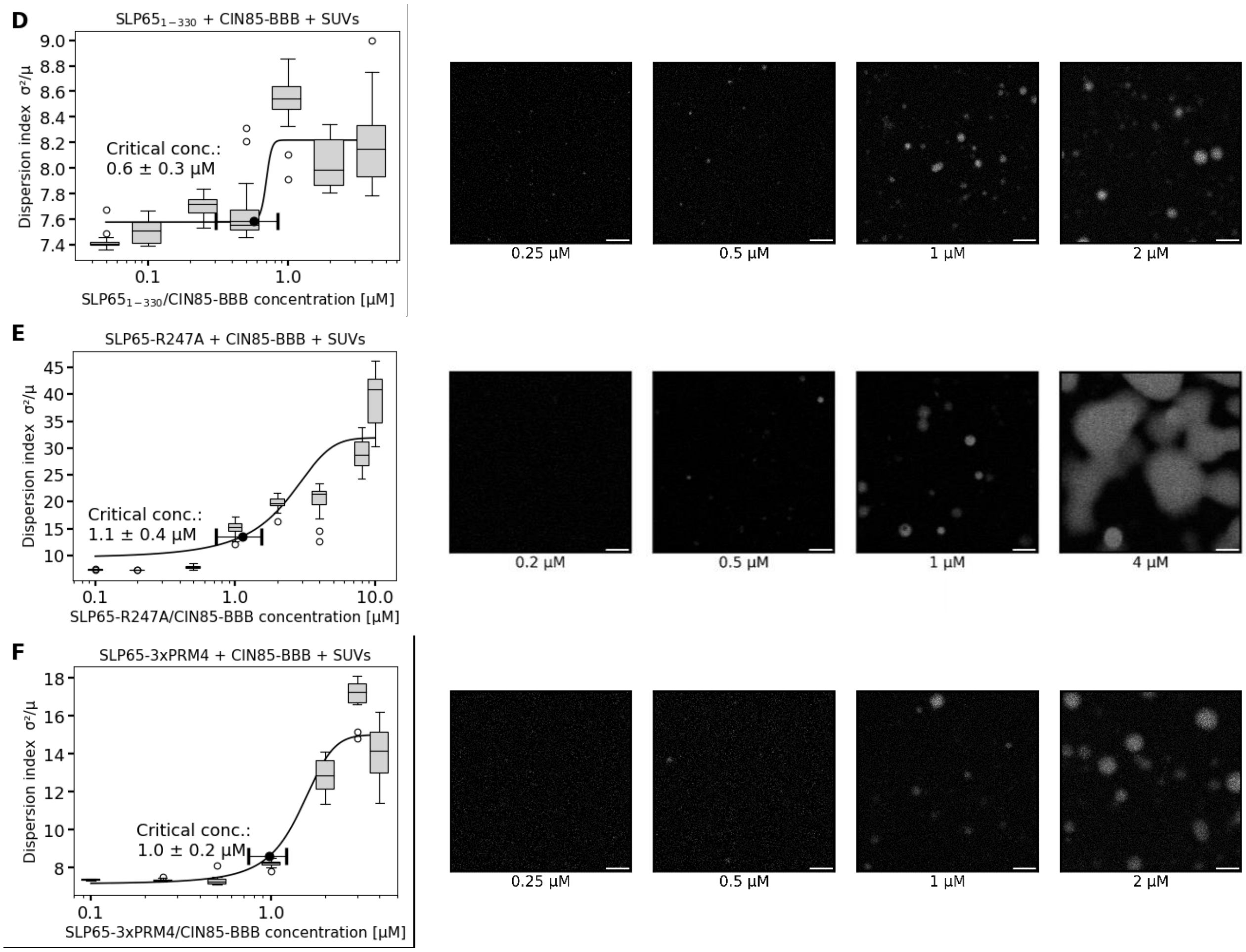


Fig. S7, continued: Critical concentrations of the CIN85-BBB construct mixed with SUVs and either SLP65$_{1-330}$ (D), SLP65-R247A (E) or SLP65-3xPRM4 (F).

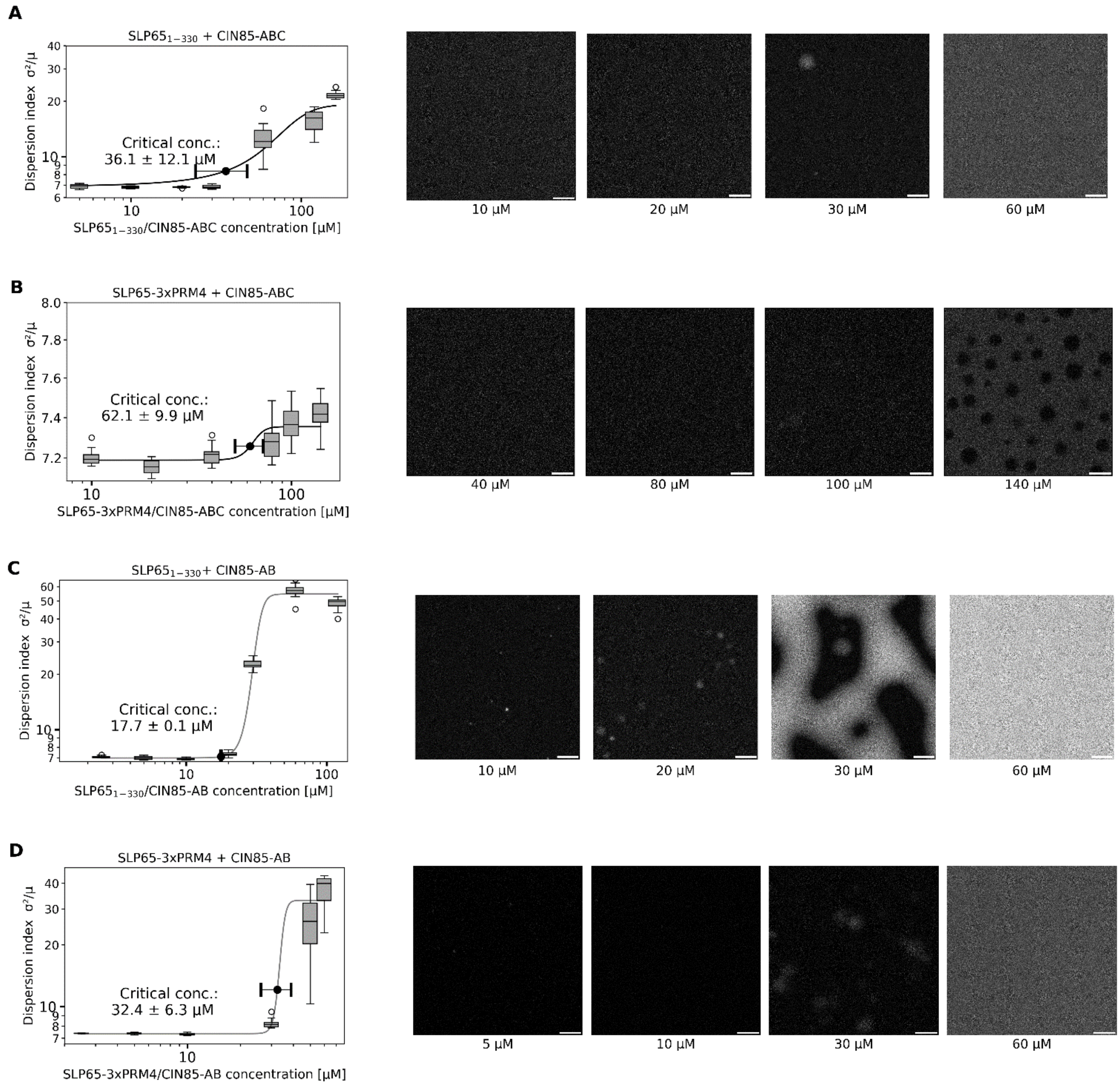


Fig. S8: Measurement of critical concentrations of SLP65/CIN85 condensates by confocal fluorescence microscopy: Two-component mixtures were prepared at equimolar SLP65/CIN85 concentrations including CIN85-ABC with either SLP65$_{1-330}$ (A) or SLP65-3xPRM4 (B), CIN85-AB with either SLP65$_{1-330}$ (C) or SLP65-3xPRM4 (D), CIN85-BBB with either SLP65-R247A (E), SLP65$_{1-330}$ (F) or SLP65-3xPRM4 (G) and CIN85-ABC-R227A-R229A with either SLP65$_{1-330}$ (H) or SLP65-3xPRM4 (I). The critical concentrations with bootstrap SD errors (highlighted in green) were obtained from fitting the dispersion index (left). Representative images (right) of mixtures show no phase separation at low concentration and phase separation at high concentrations. The mixtures including the CIN85-ABC or the CIN85-AB construct (A-D) phase separate at larger critical concentrations than the mixtures containing the CIN85-BBB construct or the CIN85-ABC-R227A-R229A construct (E-I).

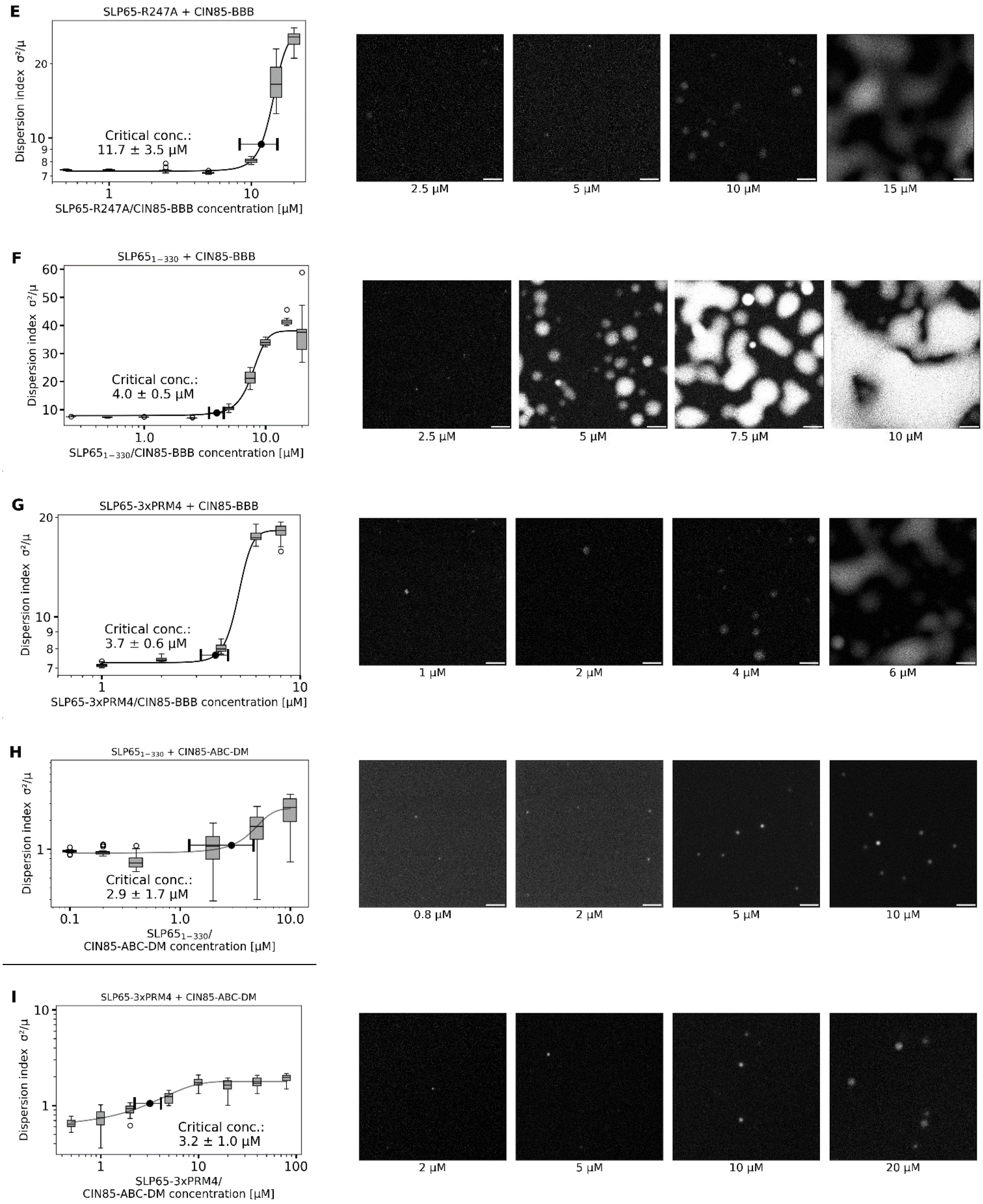


Fig. S 8 (continued): Supplementary Fig. 8:

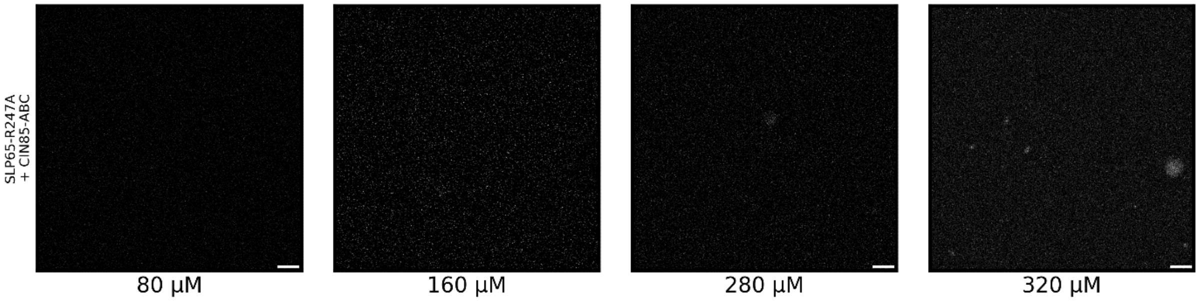


Fig. S9: Confocal microscopy images of the SLP65-R247A construct mixed with the CIN85-ABC construct. The dispersion index could not be fitted, even though a few droplets were observed at 320 µM equimolar SLP65/CIN85 concentration. Thus, no critical concentration was obtained in our measurements. Since no droplets are observed up to an equimolar concentration 280 µM $\varphi_{exp}$ is > 280 µM.

- Energy terms of the isotropic pairwise interaction energy matrix were set to 0.
- Energy terms E of the anisotropic pairwise interaction energy matrix were set equal to the $\Delta G$, which were derived from the binding constant:

| $K_D$ [μM] | PRM1 | PRM2 | PRM3 | PRM4 | PRM5 | PRM6 | SH3A | SH3B | SH3C |
|---|---|---|---|---|---|---|---|---|---|
| PRM1 | 0 | 0 | 0 | 0 | 0 | 0 | 175 | 241 | 630 |
| PRM2 | ... | 0 | 0 | 0 | 0 | 0 | 3770 | 2712 | 4169 |
| PRM3 | ... | ... | 0 | 0 | 0 | 0 | 342 | 501 | 3036 |
| PRM4 | ... | ... | ... | 0 | 0 | 0 | 159 | 6 | 35 |
| PRM5 | ... | ... | ... | ... | 0 | 0 | 107 | 54 | 209 |
| PRM6 | ... | ... | ... | ... | ... | 0 | 149 | 96 | 89 |
| SH3A | ... | ... | ... | ... | ... | ... | 0 | 0 | 0 |
| SH3B | ... | ... | ... | ... | ... | ... | ... | 0 | 0 |
| SH3C | ... | ... | ... | ... | ... | ... | ... | ... | 0 |

$$\Delta G = RT\ln(K)$$

$\Delta G$ in units of [kCal/mol]

NMR experiments at 37 °C

$$\Delta G = E$$

Anisotropic pairwise interaction energy matrix:

| #SC_SC_POT | PRM1 | PRM2 | PRM3 | PRM4 | PRM5 | PRM6 | SH3A | SH3B | SH3C |
|---|---|---|---|---|---|---|---|---|---|
| PRM1 | 0 | 0 | 0 | 0 | 0 | 0 | -5.33 | -5.13 | -4.54 |
| PRM2 | ... | 0 | 0 | 0 | 0 | 0 | -3.44 | -3.64 | -3.38 |
| PRM3 | ... | ... | 0 | 0 | 0 | 0 | -4.92 | -4.68 | -3.57 |
| PRM4 | ... | ... | ... | 0 | 0 | 0 | -5.39 | -7.38 | -6.32 |
| PRM5 | ... | ... | ... | ... | 0 | 0 | -5.63 | -6.06 | -5.22 |
| PRM6 | ... | ... | ... | ... | ... | 0 | -5.43 | -5.70 | -5.74 |
| SH3A | ... | ... | ... | ... | ... | ... | 0 | 0 | 0 |
| SH3B | ... | ... | ... | ... | ... | ... | ... | 0 | 0 |
| SH3C | ... | ... | ... | ... | ... | ... | ... | ... | 0 |

Fig. S10**:** Illustration of the symmetric, pairwise anisotropic interaction matrices. A value of the matrix refers to the interaction energy of a distinct sticker-sticker pair. The terms define the interaction energy E, which is used to calculate the interaction's equilibrium probability $p \propto \exp(-\beta * k_B * E)$, where β is the inverse simulation temperature MC_Temp, and the Boltzmann constant is defined as $k_B$ = 1 energy units / temperature units. The matrix terms were experimentally parametrized by the Gibbs free energy ΔG. Sticker-Sticker interactions of either PRM with PRM or SH3 domain with SH3 domain were set to zero. The isotropic interaction energy terms were set to 0 to allow for simulations with short linker lengths.

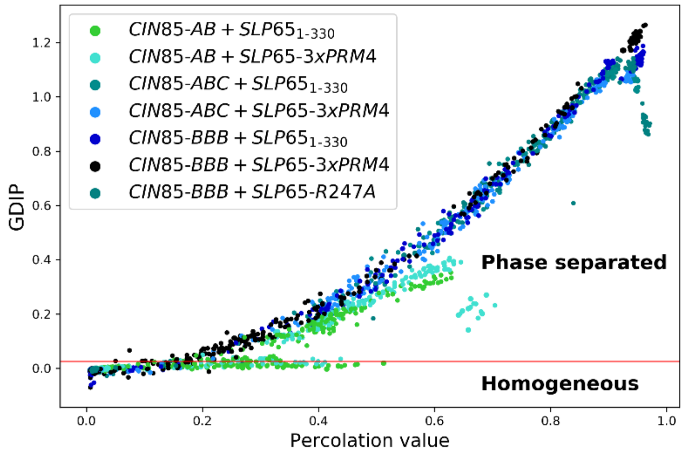


Fig. S11: Percolation value vs GDIP correlation plot. The global density inhomogeneity parameter (GDIP) scales with the percolation value except for some simulations of *CIN85-AB* with either *SLP65*$_{1\text{-}330}$ or *SLP65-3xPRM4*. The GDIP threshold for phase separation = 0.025 is shown as red line.

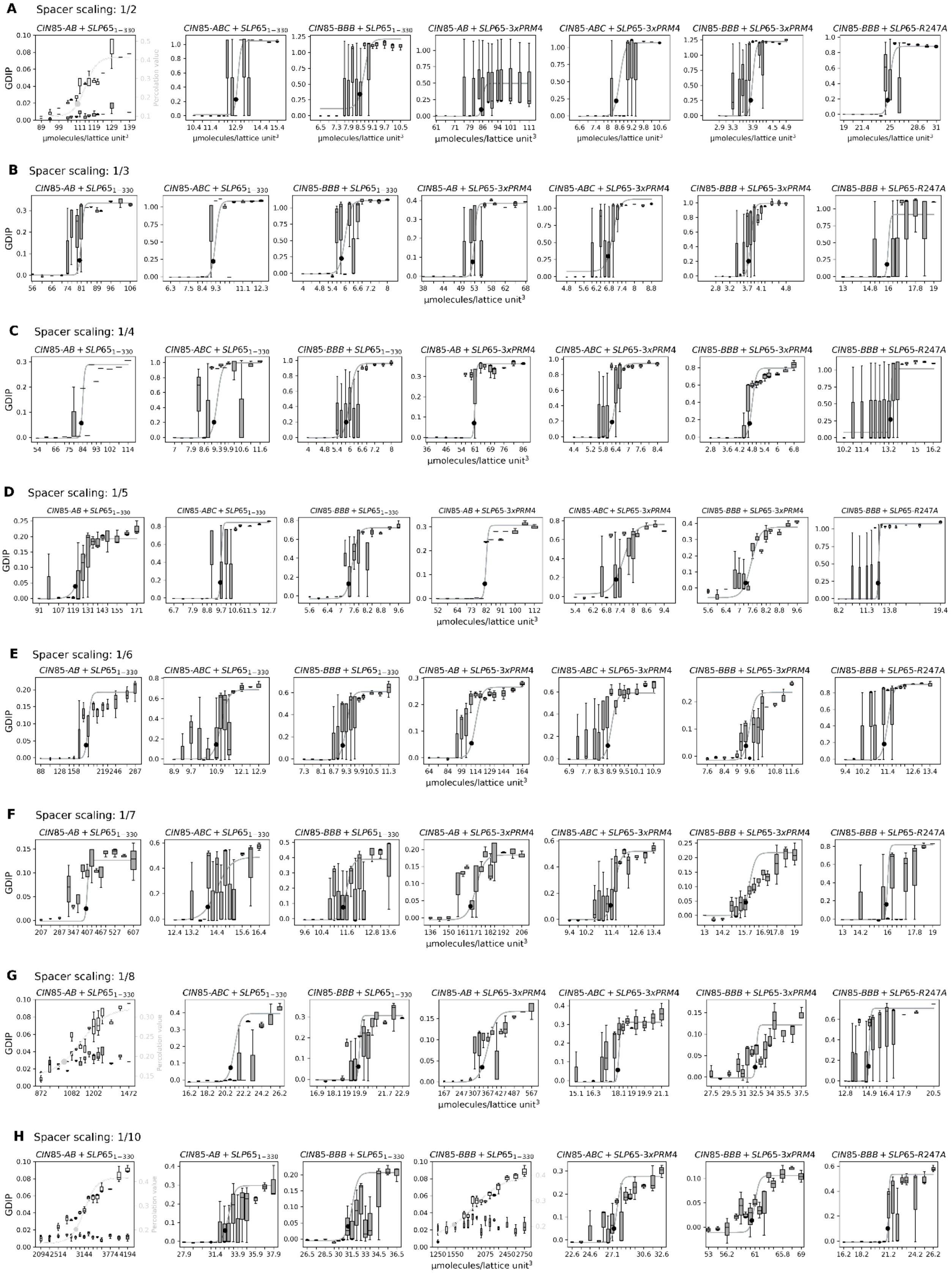

A Spacer scaling: 1/2
CIN85-AB + SLP65$_{1-330}$
CIN85-ABC + SLP65$_{1-330}$
CIN85-BBB + SLP65$_{1-330}$
CIN85-AB + SLP65-3xPRM4
CIN85-ABC + SLP65-3xPRM4
CIN85-BBB + SLP65-3xPRM4
CIN85-BBB + SLP65-R247A
B Spacer scaling: 1/3
C Spacer scaling: 1/4
D Spacer scaling: 1/5
E Spacer scaling: 1/6
F Spacer scaling: 1/7
G Spacer scaling: 1/8
H Spacer scaling: 1/10
CIN85-BBB + SLP65$_{1-330}$
GDIP
Percolation value
μmolecules/lattice unit$^3$

Fig. S12: Simulations of the critical concentration of two-component mixtures modeling the different-affine designer constructs either *SLP651-330* or *SLP65-PRM4* with either *CIN85-AB*, *CIN85-ABC* or *CIN85-BBB*. The spacer were varied by applying a scaling factor between 1/2 and 1/10 (A-H). In case were the GDIP was too low, the percolation value was fitted, which scales with the GDIP (Fig. S11).

| Simulated combination | Spacer scaling: 1/2 | 1/3 | 1/4 | 1/5 | 1/6 | 1/7 | 1/8 | 1/10 | |
|---|---|---|---|---|---|---|---|---|---|
| | $\phi_{sim}$ [µmolecules/lattice-unit$^3$] | | | | | | | | |
| *CIN85-AB + SLP65$_{1-330}$* | 110* | 80 | 83 | 122 | 184 | 402 | 1030* | 2941* | **Large ϕ** |
| *CIN85-AB + SLP65-3xPRM4* | 85 | 52 | 60 | 81 | 110 | 167 | 341 | 1529* | |
| *CIN85-ABC + SLP65$_{1-330}$* | 12.9 | 9.1 | 9.1 | 9.5 | 10.9 | 14 | 21 | 32 | |
| *CIN85-ABC + SLP65-3xPRM4* | 8.5 | 6.8 | 6.2 | 7.2 | 9.5 | 11.3 | 18 | 27 | |
| *CIN85-BBB + SLP65$_{1-330}$* | 8.4 | 5.8 | 5.8 | 7.4 | 9.1 | 11.4 | 20 | 31 | |
| *CIN85-BBB + SLP65-3xPRM4* | 3.8 | 3.7 | 4.7 | 7.3 | 9.4 | 15.7 | 32 | 60 | |
| *CIN85-BBB + SLP65-R247A* | 24.7 | 15.9 | 13.3 | 12.5 | 11.2 | 16 | 15 | 21 | **Small ϕ** |

Fig. S13: Simulated critical concentrations (φsim) of combinations of the CIN85-AB, CIN85-ABC or CIN85-BBB with SLP651-330, SLP65-3xPRM4 or SLP65-R247A molecules. The lengths of the spacer were varied by scaling the natural linker length between 1/2 and 1/10

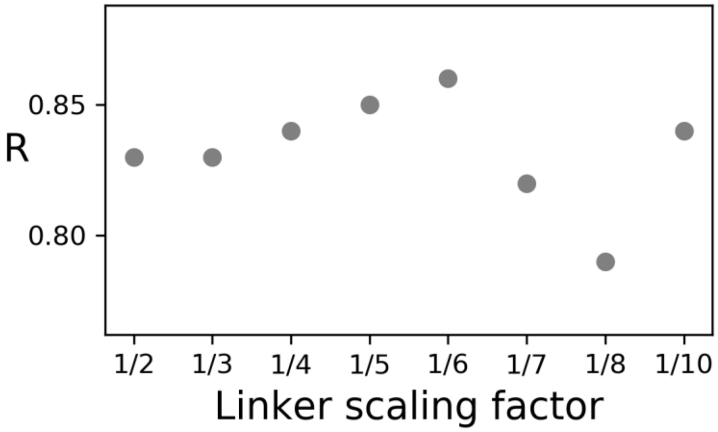


Fig. S14: Pearson correlation coefficients for linker length scaling factors between ½ - 1/10. The Pearson correlation coefficient is calculated from nine pairs of $\varphi_{exp}$- *vs* $\varphi_{sim}$-values, excluding the correlations of CIN85-ABC + SLP65$_{1-330}$/SLP65-3xPRM4 with *CIN85-ABC* + *SLP65$_{1-330}$/SLP65-3xPRM4* (see Fig. 4).

Table S1: ITC binding constants and thermodynamic parameters from titrations of the designer constructs of SLP65 with CIN85.

| Titration | #Repli-cates | N | $K_D$ [µM] | ΔH [kcal/mol] | ΔS [kcal/mol*K] |
|---|---|---|---|---|---|
| A) $SLP65_{1\text{-}330}$ to CIN85-ABC | 3 | 1.15 ± 0.07 | 1.36 ± 0.17 | -19.56 ± 0.69 | -36.3 |
| B) SLP65-3xPRM4 to CIN85-ABC | 3 | 0.75 ± 0.07 | 0.24 ± 0.01 | -24.66 ± 0.2 | -49.3 |
| C) SLP65-R247A to CIN85-ABC | 3 | 2.72 ± 0.79 | 7.04 ± 1.12 | -11.77 ± 1.97 | -14.5 |
| D) $SLP65_{1\text{-}330}$ to CIN85-BBB | 3 | 1.74 ± 0.44 | 0.7 ± 0.25 | -5.03 ± 0.76 | 11.6 |
| E) SLP65-3xPRM4 to CIN85-BBB | 2 | 0.78 ± 0.08 | 0.14 ± 0.06 | -15.68 ± 2.08 | -19.3 |
| F) SLP65-R247A to CIN85-BBB | 3 | 5.77 ± 2.16 | 16.1 ± 8.56 | -3.51 ± 2.33 | 10.4 |

Table S2: Parameter settings for the CIN85 and SLP65 linkers. In the one-bead per domain/motif LASSI model, spacers are defined by scaling the natural linker length.

**Spacers** [lattice units]

| | Amino acids | Scaling factors | | | | | | | |
|---|---|---|---|---|---|---|---|---|---|
| | | 1/2 | 1/3 | 1/4 | 1/5 | 1/6 | 1/7 | 1/8 | 1/10 |
| $CIN85_{1\text{-}333}$: | | | | | | | | | |
| **Linker 1** | 39 | 20 | 13 | 10 | 8 | 7 | 6 | 5 | 4 |
| **Linker 2** | 66 | 33 | 22 | 16 | 13 | 11 | 9 | 8 | 7 |
| **Linker 3** | 38 | 19 | 13 | 10 | 8 | 6 | 5 | 5 | 4 |
| $SLP65_{1\text{-}330}$: | | | | | | | | | |
| **Linker 1** | 60 | 30 | 20 | 15 | 12 | 10 | 9 | 8 | 6 |
| **Linker 2** | 45 | 23 | 15 | 11 | 9 | 8 | 6 | 6 | 5 |
| **Linker 3** | 74 | 37 | 25 | 19 | 15 | 12 | 11 | 9 | 7 |
| **Linker 4** | 27 | 14 | 9 | 7 | 5 | 5 | 4 | 3 | 3 |
| **Linker 5** | 26 | 13 | 9 | 7 | 5 | 4 | 4 | 3 | 3 |

Table S3: NMR titration experiments of SLP65- and CIN85-derived peptides (ligand) to the SH3A, SH3B or SH3C domain (receptor) was performed at the indicated receptor concentration and ligand/protein ratios.

| **Ligand: Peptide X** | **Receptor: CIN85-SH3 domain** | **Receptor concentration [µM]** | **Molar ligand/receptor ratios** |
|---|---|---|---|
| SLP65-PRM1 | SH3A | 200 | 0.2, 0.5, 0.7 , 0.9, 1.1, 1.4 , 1.6, 1.8, 2.0, 2.6, 3.3 |
| SLP65-PRM2 | SH3A | 200 | 0.6, 1.2, 1.8, 2.4, 3.2, 4.7 |
| SLP65-PRM3 | SH3A | 200 | 0.5, 0.9, 1.4, 1.8, 2.3, 2.7, 3.5, 4.3 |
| SLP65-PRM4 | SH3A | 200 | 0.3, 0.6, 0.9, 1.2, 1.5, 2.4, 4.1, 7.1, 10.2, 10.7 |
| SLP65-PRM5 | SH3A | 200 | 0.5, 0.9, 1.5, 1.8, 1.9, 2.2, 2.6, 3.8, 5.7, 9.0 |
| SLP65-PRM6 | SH3A | 200 | 0.3, 0.6, 0.9, 1.2, 1.5, 2.4, 3.3, 4.7, 9.0 |
| SLP65-PRM1 | SH3B | 260 | 0.2, 0.5, 0.8, 1.1, 1.3, 1.6, 2.2, 2.8, 3.4, 4.0 |
| SLP65-PRM2 | SH3B | 200 | 0.3, 0.6, 0.9, 1.2, 1.8, 2.7, 4.7, 7.4, 10.0 |
| SLP65-PRM3 | SH3B | 260 | 0.5, 0.9, 1.4, 1.8, 2.3, 2.7, 3.5, 4.3, 5.6 |
| SLP65-PRM4 | SH3B | 200 | 0.6, 1.3, 1.6, 2.5, 3.4, 4.1, 5.0, 10.0 |
| SLP65-PRM5 | SH3B | 200 | 0.3, 0.9, 1.2, 1.5, 2.1, 3.0, 3.4, 4.2, 8.0 |
| SLP65-PRM6 | SH3B | 200 | 0.6, 0.9, 1.2, 1.5, 2.4, 3.3, 4.7 |
| SLP65-PRM1 | SH3C | 200 | 0.3, 0.6, 0.9, 1.2, 1.5, 2.4, 3.3, 4.7, 9.0 |
| SLP65-PRM2 | SH3C | 200 | 0.6, 1.2, 1.8, 2.4, 3.3, 4.7, 8.9, 12.8 |
| SLP65-PRM3 | SH3C | 200 | 0.9, 1.2, 1.5, 2.4, 3.3, 4.7, 9.0 |
| SLP65-PRM4 | SH3C | 200 | 0.3, 0.6, 0.9, 1.2, 1.5, 2.4, 3.3, 4.7, 9.0 |
| SLP65-PRM5 | SH3C | 200 | 0.3, 0.6, 0.9, 1.2, 1.5, 2.4, 3.3, 4.7, 9.0 |
| SLP65-PRM6 | SH3C | 200 | 0.9, 1.2, 1.5, 2.4, 3.3, 4.7, 9.0 |

Table S4: Injection scheme for ITC titrations on the VP-ITC MicroCalorimeter (MicroCal). SLP65-peptides derived from PRM1 to PRM6 were titrated to SH3A, SH3B and SH3C domains. $SLP65_{1-330}$ and weak- and strong-binding SLP65 constructs were titrated to CIN85-ABC/BBB constructs. Total injection volume was 284 µl. Binding isotherms of peptide titrations were fitted with the single-site model (Origin 7) fixing the n-value = 1 (no floating n value). Binding isotherms of multivalent protein constructs were fitted with the single-site model (Origin 7) due to the lack of complete data set for global analysis [13].

| Injection number | Injection volume [µl] | Duration of injection [ms] | Time between injection points: Spacing [s] | Filter Period [s] |
|---|---|---|---|---|
| 1 | 4.0 | 8.0 | 400 / 600 | 2 |
| 2 | 20.0 | 40.0 | 400 / 600 | 2 |
| 3 | 20.0 | 40.0 | 400 / 600 | 2 |
| 4 | 20.0 | 40.0 | 400 / 600 | 2 |
| 5 | 20.0 | 40.0 | 400 / 600 | 2 |
| 6 | 20.0 | 40.0 | 400 / 600 | 2 |
| 7 | 20.0 | 40.0 | 400 / 600 | 2 |
| 8 | 20.0 | 40.0 | 400 / 600 | 2 |
| 9 | 20.0 | 40.0 | 400 / 600 | 2 |
| 10 | 20.0 | 40.0 | 400 / 600 | 2 |
| 11 | 20.0 | 40.0 | 400 / 600 | 2 |
| 12 | 20.0 | 40.0 | 400 / 600 | 2 |
| 13 | 20.0 | 40.0 | 400 / 600 | 2 |
| 14 | 20.0 | 40.0 | 400 / 600 | 2 |
| 15 | 20.0 | 40.0 | 400 / 600 | 2 |
